# Benchmarking UMI-based single cell RNA-sequencing preprocessing workflows

**DOI:** 10.1101/2021.06.17.448895

**Authors:** Yue You, Luyi Tian, Shian Su, Xueyi Dong, Jafar S Jabbari, Peter F Hickey, Matthew E Ritchie

## Abstract

Single-cell RNA sequencing (scRNA-seq) technologies and associated analysis methods have undergone rapid development in recent years. This includes methods for data preprocessing, which assign sequencing reads to genes to create count matrices for downstream analysis. Several packaged preprocessing workflows have been developed that aim to provide users with convenient tools for handling this process. How different preprocessing workflows compare to one another and influence downstream analysis has been less well studied.

Here, we systematically benchmark the performance of 9 end-to-end preprocessing workflows (*Cell Ranger, Optimus, salmon alevin, kallisto bustools, dropSeqPipe, scPipe, zUMIs, celseq2* and *scruff*) using datasets with varying levels of biological complexity generated on the CEL-Seq2 and 10x Chromium platforms. We compare these workflows in terms of their quantification properties directly and their impact on normalization and clustering by evaluating the performance of different method combinations. We find that lowly expressed genes are discordant between workflows and observe that some workflows have systematic biases towards particular classes of genomics features. While the scRNA-seq preprocessing workflows compared varied in their detection and quantification of genes across datasets, after downstream analysis with performant normalization and clustering methods, almost all combinations produced clustering results that agreed well with the known cell type labels that provided the ground truth in our analysis.

In summary, the choice of preprocessing method was found to be less influential than other steps in the scRNA-seq analysis process. Our study comprehensively compares common scRNA-seq preprocessing workflows and summarizes their characteristics to guide workflow users.

## Background

Over the past decade, single-cell RNA sequencing (scRNA-seq) technologies and associated analysis methods have rapidly developed and been applied to a wide range of biological systems (1, 2). The large number of analysis methods available presents a significant challenge for data analysts who are left to choose which of the many tools are best suited to their experiment and analysis goals. Fortunately, there are now many systematic benchmarking studies that explore this question in detail (3–6). However, these evaluations almost exclusively focus on downstream analysis tasks, including normalization, clustering, trajectory analysis, cell type identification and data integration. They ignore the crucial first step of preprocessing that summarises the sequencing reads into a count matrix which is used as input to all downstream analyses.

The main difference between preprocessing scRNA-seq data compared with bulk RNA-seq lies in having to deal with various DNA barcodes which assist in the assignment of sequence reads to their cell or molecule of origin (7, 8). Most methods are designed to work with unique molecular identifier (UMI) (9, 10) based data, since these protocols are widely used in the field. Typical scRNA-seq preprocessing workflows involve demultiplexing, mapping, transcript quantification and quality control. An overview of the main steps involved in preprocessing is summarised in Figure 1A. Starting from raw `FASTQ` files, cell barcodes (CBs) and UMIs are first appended as tags to the header of each cDNA read. Next, cDNAs are aligned to either a reference genome or transcriptome or pseudoaligned to the transcriptome depending on the particular pipeline. To overcome amplification bias, UMIs are used to remove PCR-duplicated molecules from the gene counts in each cell. Base errors in CBs and UMIs are usually corrected at this step or before alignment. Next, reads are separated by CBs and assigned to genes or transcripts which allows construction of a cell-by-gene count matrix (with cells in the columns and genes/transcripts in rows). Next, cells of low quality and genes with low abundance are typically filtered out and the resulting count matrix is used in down-stream analysis. Of note, CBs are known in advance for each well in plate-based protocols such as CEL-Seq (11) and randomly assigned to cells in droplet-based protocols, like 10x Chromium (12) and inDrops (13). Because of this, different strategies are applied to construct CB ‘allow lists’ and additional steps to distinguish real cells from ambient RNA are suggested for droplet-based protocols (14).

**Fig. 1.**
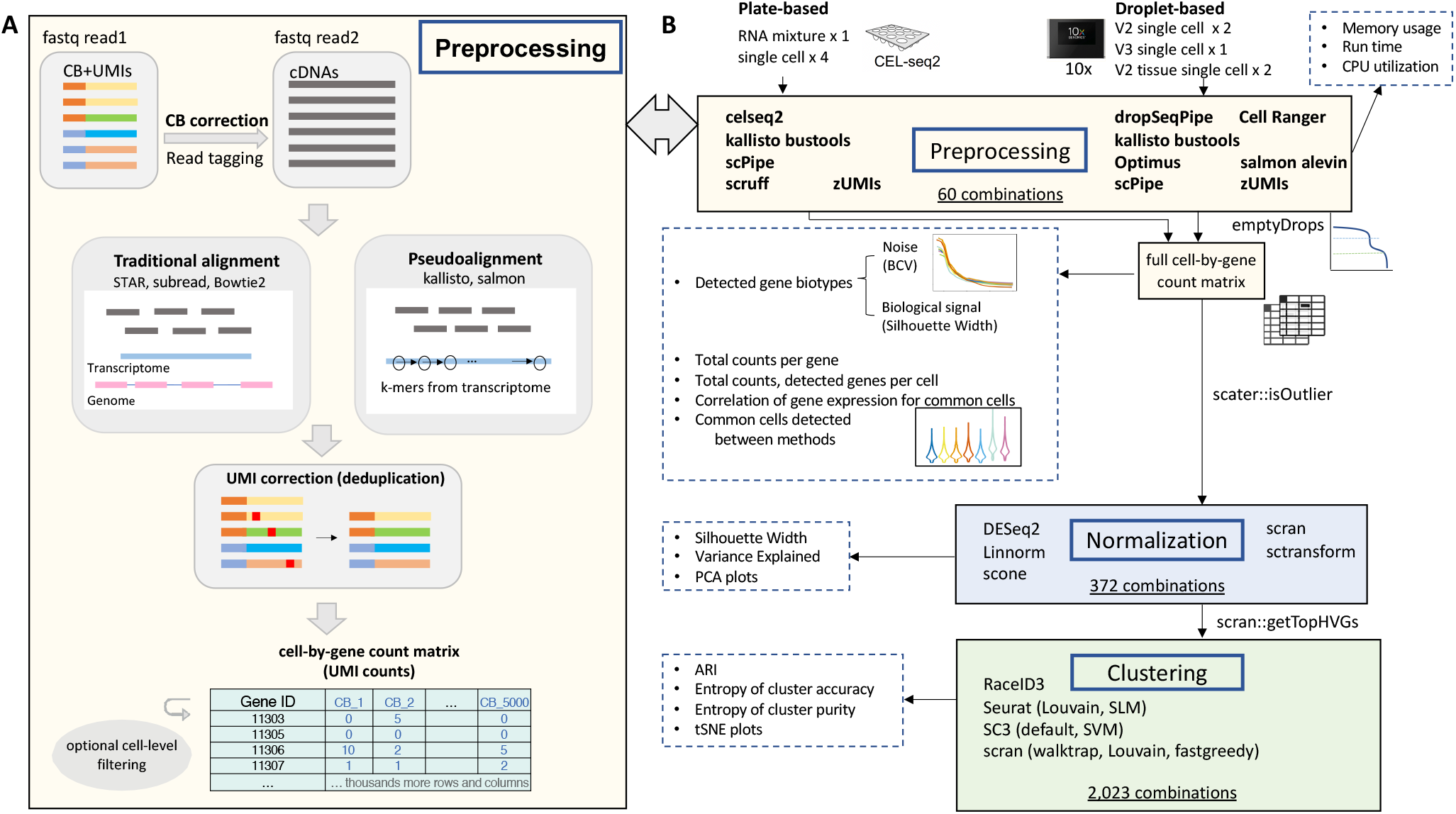
Overview of scRNA-seq preprocessing workflows and our benchmarking study design. **A)** A typical preprocessing workflow begins with raw sequences in `FASTQ` files, that are subject to cell barcode (CB) detection, alignment, UMI correction, count matrix generation and quality control. **B)** Summary of our benchmarking study, showing the datasets analyzed, the selected preprocessing workflows, methods for normalization and clustering that were compared. Workflows and methods used in analysis are listed in boxes with solid borders, while evaluation metrics are shown in boxes with dashed borders. In total, 2,023 combinations of datasets × preprocessing workflows × downstream analysis methods were generated in this benchmarking study.

Researchers can build their own preprocessing workflows by combining individual methods that address each of the aforementioned steps, or they may choose one of the many packaged workflows that have been proposed and published that aim to make this process more convenient. Examples of preprocessing workflows include *Cell Ranger* (12), *UMI-tools* (15), *scPipe* (16) and *zUMIs* (17). More recently, alignment-free tools such as *salmon* (18) and *kallisto* (19) have been adapted to handle single-cell data to improve the computational efficiency of scRNA-seq data analysis. In addition, projects like the Human Cell Atlas (20) have developed their own preprocessing workflow (*Optimus* (21)) that uniformly processes the millions of human single-cell transcriptomes generated through this international collaboration. Other examples include the Single Cell Expression Atlas that has applied the *SCXA* pipeline (https://github.com/ebi-gene-expression-group/scxa-workflows) to process in excess of 4 million cells to build a large cross-species collection of single cell expression profiles (22).

Differences between published workflows arise when researchers balance efficiency with accuracy at each of the above-mentioned steps. For example, to obtain CBs, most workflows use a ‘allow list’ as a reference, whereas *salmon alevin* generate a putative list of highly abundant CBs that can be further filtered. To deduplicate UMIs, *kallisto bustools* (23) applied a naive collapsing strategy that they found to be more effective than more complicated approaches.

Other pipelines such as *Cell Ranger* and *zUMIs* also take base quality and edit distance into consideration during UMI deduplication. Methods that place more importance on this step have also been developed after finding edit distances were inadequate for dealing with UMIs with high similarity (24, 25). For example, *UMI-tools* introduced a network-based graph approach for this step, *alevin* (24) constructed parsimonious UMI graphs and *dropEst* (25) developed a Bayesian method to model UMI errors. To assign multi-mapped reads, several workflows, including *Cell Ranger, Optimus, dropSeqPipe* and *kallisto bustools* discard them, while others treat ambiguous reads in different ways, assigning them to potential mapping positions probabilistically or via other strategies.

A small number of studies have compared the performance of different scRNA-seq preprocessing algorithms and they tend to stop at the point before normalization after observing high correlations between count matrices obtained from the above-mentioned workflows or other custom combinations of methods for each step (17, 26). Quantification differences were studied by comparing the performance of various popular alignment methods and annotation schemes (27) to help guide the choice of such tools. Based on present preprocessing workflows, one study compared the performance of high-throughput scRNA-seq pipelines before and after normalization followed by clustering and differential expression analysis (28). They pointed out a confounding factor akin to batch effects after integrating matrices processed by multiple different workflows applied to the same dataset. However, they did not utilize datasets with gold standard ground truth, and pseudo-alignment based workflows such as *kallisto bustools* and *salmon alevin* were not included in the comparison. A more recent study compared the performance of these two pseudo-alignment pipelines, and demonstrated that *kallisto bustools* was faster and used less memory while producing similar downstream results (29). In contrast, others found *kallisto bustools* results to have an overrepresentation of cells with low gene content which are likely due to mapping artifacts (30). Preprocessing tools have also been shown to influence the results of RNA velocity analyses (31) which highlights the potential for downstream effects driven by pre-processing algorithm choice. It is worth noting that previous comparative studies mostly focus on preprocessing for droplet-based protocols thus ignoring plate-based platforms, like CEL-Seq (11) and CEL-Seq2 (32) which are frequently used in some settings.

Here, we systematically benchmarked 9 end-to-end preprocessing workflows, including *scPipe, zUMIs, kallisto bustools, dropSeqPipe* (33), *Cell Ranger, Optimus, salmon alevin, celseq2* (32) and *scruff* (34). Among them, *celseq2* and *scruff* are specific to data generated by CEL-Seq and CEL-Seq2 protocols. *Cell Ranger* was developed for use with the 10x Chromium platform and is the standard workflow for 10x datasets. *DropSeqPipe* is only available for droplet-based protocols and is an unpublished online workflow with an instructional video on YouTube that teaches users how to run it. *Alevin* is a tool integrated within *salmon* that proposes new methods to handle UMIs and ambiguous reads. *scPipe, zUMIs, salmon alevin* and *kallisto bustools* can all handle raw data from both plate and droplet-based platforms, and the first three can also deal with Smart-Seq (35) (a full-length protocol that is not UMI-based) data.

We apply these methods to various scRNA-seq datasets with available ground truth containing varying biological complexity levels to benchmark their performance. Specifically, we describe the basic features of the count matrix produced by each preprocessing workflow and explore the impact on downstream analysis by evaluating the performance of combinations of preprocessing workflows together with various normalization and clustering methods using the *CellBench* platform (36).

## Results

### Benchmarking scRNA-seq preprocessing workflows

#### scRNA-seq preprocessing workflows evaluated

We investigated the performance of 9 end-to-end workflows applied to plate-based (CEL-Seq2) and droplet-based (10x Chromium v2 and v3 chemistry) data. To process CEL-Seq2 data, we applied *celseq2* and *scruff* (only applicable to plate-based protocols) along with *scPipe, zUMIs*, and *kallisto bustools* (applicable to both plate- and droplet-based protocols). For the 10x data we applied *dropSeqPipe, Cell Ranger, Optimus, salmon alevin, scPipe, zUMIs* and *kallisto bustools*. Details of each workflow and the specific strategies applied within preprocessing are listed in Table S1.

#### scRNA-seq datasets used for benchmarking

The *scmixology* datasets (5), which were designed for scRNA-seq bench-marking studies, include cells from distinct cell lines and provide ground truth in various forms (e.g., known clusters based on the mixing strategy applied or based on genetic variation between cell lines). These datasets involve experimental designs that use controlled mixtures of RNA (*RNA mixture*, 1 × 384-well plate of CEL-Seq2 data) to create ‘pseudo-cells’, or actual single cells from up to 5 human lung adenocarcinoma cell lines (4 × 384-well plates of CEL-Seq2 data and 2 × 10x Chromium v2 datasets). A new dataset using the same five cell lines profiled with 10x Chromium v3 chemistry was also generated (data available from GEO under accession number GSE154870). The final datasets included in our analysis were from the Tabula Muris project (37). Cells from mouse lung tissue profiled by 10x Chromium (2 × 10x Chromium v2 datasets) were included to assess performance in a setting with more cellular diversity than the *scmixology* datasets.

The datasets varied in terms of their experimental design, cell number, and the level of biological noise as summarized below:

- ‘low’ noise setting: *pseudo-cells* created by mixing RNA in 7 different concentrations (groups).
- ‘medium’ noise setting: single cells obtained from cell lines (3 or 5 different groups).
- ‘high’ noise setting: single cells obtained from tissue samples assigned to 13 different clusters.

A summary of the datasets used, including the sample size, expected number of clusters, and data structure is given in Table S2.

Cell labels provided by the *scmixology* datasets were generated with intermediate BAM files created by *scPipe* based on single-nucleotide polymorphisms (SNPs) information for the single cell datasets and via labels available from the plate annotation for the mixture experiments. Cells from the Tabula Muris samples were manually annotated using canonical marker genes according to the approach described in Tabula Muris Consortium (2018) (37), and these cell type labels were used as the ground truth in our study.

### Benchmarking workflow

An overview of our benchmarking study design is presented in Figure 1B. We generated a cell-by-gene count matrix using each of the preprocessing workflows listed in Table S1. We performed cell-level quality control by firstly applying *emptyDrops* to distinguish empty droplets and cells (droplet-based protocols) and then using *scater* to identify and remove low-quality cells by setting a data-driven threshold on various quality control metrics (applied to both plate- and droplet-based protocols). Count matrices were normalized by five representative normalization methods that included *scran, Linnorm, scone, DESeq2* and *sctransform*. We then selected highly variable genes (HVGs) and applied up to eight commonly used clustering methods, including *RaceID3, Seurat* with the smart local moving algorithm (SLM) and Louvain algorithms, *SC3, SC3* with the SVM and *scran* with the walktrap, Louvain and fastgreedy algorithms.

Overall, 2,023 combinations of datasets × preprocessing workflows × downstream analysis methods were obtained, with performance evaluated by several metrics at each step (see Methods and Figure 1B). *CellBench* was used to compare these combinations at the pipeline-level, which allowed us to assess both the performance of a single method at a specific processing step and the interaction of multiple methods across several steps.

### Comparing computational performance of scRNA-seq preprocessing workflows

One important consideration when choosing between preprocessing workflows is their requirement of time and memory. We compared workflows by specifying one node and eight threads on a High-Performance Computing (HPC) system. Specific memory and the amount of time is required in each submission. To control for competing workloads on the HPC, we ran each workflow with datasets of different sizes three times per dataset and recorded run time, maximum memory usage, and CPU utilization (see Methods for details). A summary of the results obtained is shown in Figure 2.

**Fig. 2.**
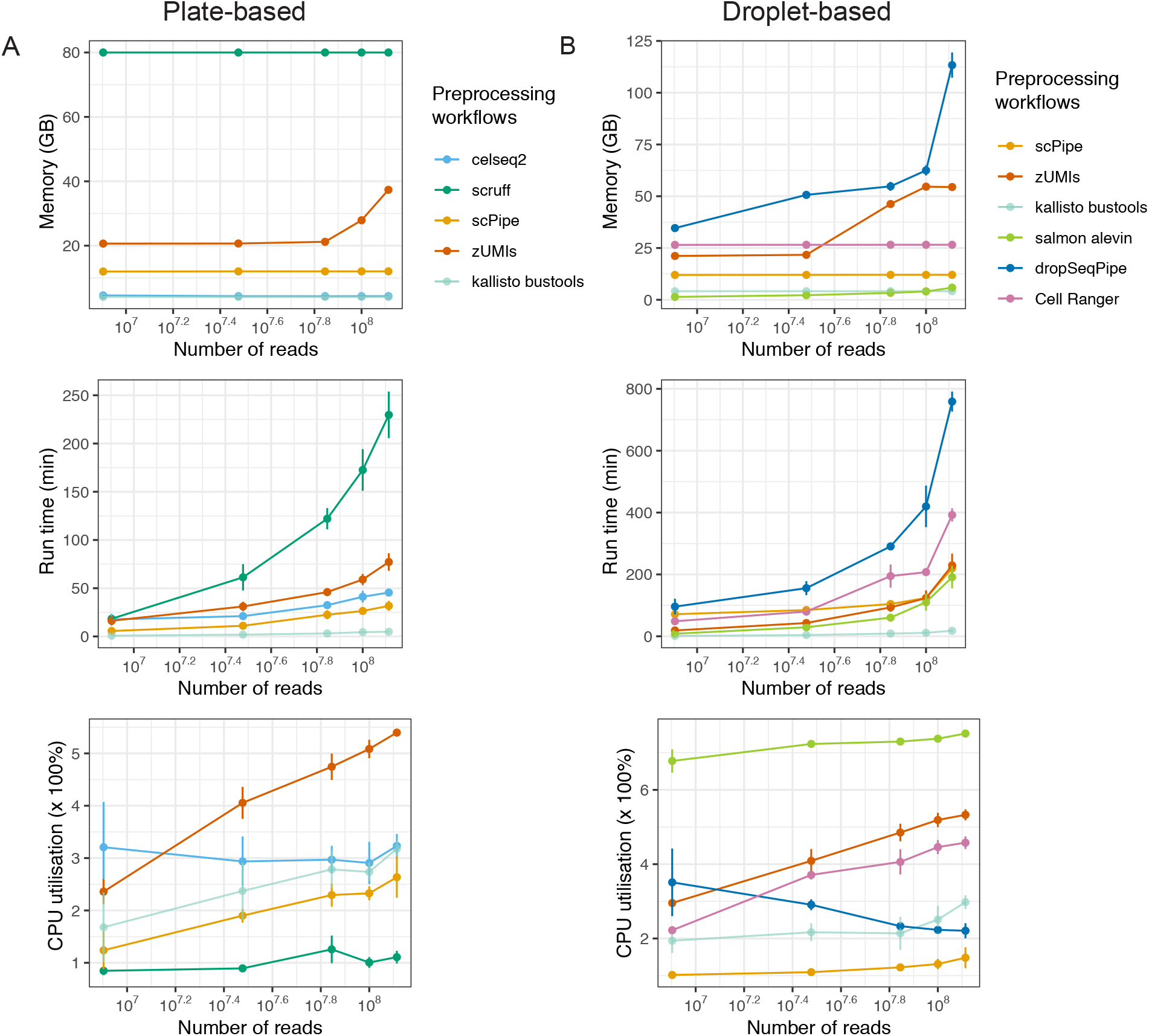
Comparing the computational performance of different scRNA-seq preprocessing workflows. CPU utilization, maximum memory usage, and run time for each preprocessing workflow. Results for preprocessing workflows designed for **A)** plate-based protocols (left-hand panel) and **B)** droplet-based protocols (right-hand panel) are shown.

For plate-based protocols, *scruff* required more memory and was slower when data volume increased, which is in contrast to the results of Wang *et al*. (2019) (34). This discrepancy might be due to their usage of smaller datasets containing fewer than 10 million reads and differences in the hardware and settings of parallelization used for evaluation. *scPipe, zUMIs* and *celseq2* showed similar maximum memory consumption and running times.

For droplet-based protocols, workflows based upon pseudo-alignment tools required less memory resource and ran faster, concordant with previous studies (18, 19). *kalliso bustools* was more than 50 times faster when dealing with 200 million reads compared to the slowest workflow *dropSeqPipe*. Maximum memory usage was at a similar level for *Cell Ranger, scPipe* and *kallisto bustools*, while other workflows used more memory as dataset size (in terms of the number of reads) increased. *salmon alevin* displayed the highest value of CPU utilization, suggesting less time is spent waiting. Another research study (28) also compared the computational performance of preprocessing workflows and demonstrated higher CPU utilization values and shorter running times for *Cell Ranger*, which is contrary to our results. This difference is likely due to our specifying a fixed number of cores and restricted memory for evaluation, whereas Gao *et al*. did not.

### Comparing gene quantification across scRNA-seq preprocessing workflows

Besides computational efficiency, the characteristics and accuracy of the biological information recovered by different methods is another key consideration when selecting a preprocessing workflow. Using all cells and genes with overall expression above zero (i.e. a count of one or more in at least one cell) without any other filtering, we characterized the workflows in terms of detected genes per cell, total counts per cell, correlation of gene expression between common cells and the concordance of retained cells identified by different workflows.

### Gene quantification for CEL-Seq2 workflows

For the CEL-Seq2 benchmarking datasets, 5 preprocessing workflows were applied: *scruff, celseq2, scPipe, zUMIs* and *kallisto bustools*. There was little variation in the different metrics assessed between CEL-Seq2 datasets, so representative results for the plate_3cl dataset are shown in Figure 3.

**Fig. 3.**
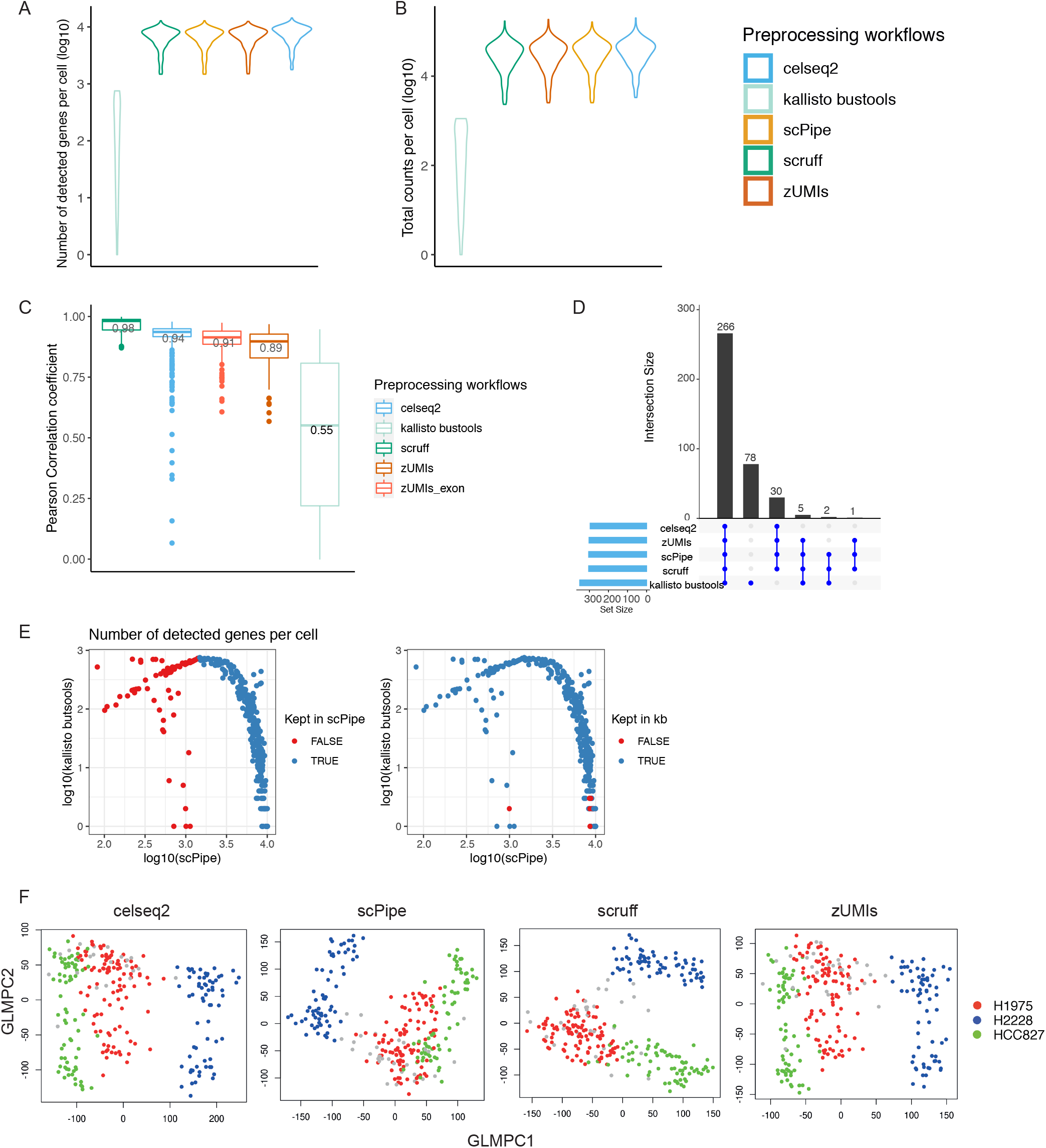
Comparing gene expression quantification of different scRNA-seq preprocessing workflows on plate_3cl dataset. **A)** Number of detected genes per cell and **B)** total counts per cell (both on a log 10-scale) are shown. The Pearson correlation coefficients of genes between *scPipe* with other preprocessing workflows are shown in **C)**. Next, after filtering, the UpSet plot in **D)** is used to display intersections of retains cells across workflows. In **E)**, number of detected genes per cell of *kallisto bustools* and *scPipe* were plotted in a pair-wise manner. Colors represent whether that cell is kept after filtering with *scPipe* in the left panel and *kallisto bustools* in the right panel. GLMPCA plots delivered across preprocessing workflows are shown in **F)**. Colors represent different cell line cells. For not common cells, cells are colored in grey.

In summary, the number of detected genes and total counts per cell showed high similarity across preprocessing workflows except for *kallisto bustools* (Figure 3A-B), which recovered fewer genes and lower counts. Upon further investigation, this was found to be related to its strategy of assigning only unique CB-UMI pairs per cell (further details on this issue can be found at https://github.com/BUStools/bustools/issues/44). For short UMIs (6 bp in the case of the CEL-Seq2 datasets), this limits the maximum detected features per cell to 46 = 4, 096 (3.61 on the log_1_ 0-scale). In practice the UMI counts are much lower, as whenever the same UMI is observed across more than one gene, it is removed from the analysis altogether. This deficiency led us to exclude *kallisto bustools* from the majority of comparison plots for the CEL-Seq2 preprocessing results.

Slightly more detected genes per cell were observed with *celseq2* (Figure 3A, S1A), suggesting higher sensitivity of the aligner it uses (*Bowtie2*) on the datasets tested. *scruff* and *scPipe* were in complete agreement concerning the number of detected genes (Figure S1A), which is as expected, as they share a very similar strategy across all preprocessing steps except for the quantification method used to count aligned reads. Higher total counts per cell were observed with *scPipe* compared with those of *scruff* (Figure S1B). We speculate that the strategy applied in *scPipe* of UMI collapsing and read quantification may give rise to fewer collapsed UMIs and more assigned reads, and consequently, higher total counts per cell.

We next investigated the concordance of gene expression across workflows and compared the correlation of gene expression between *scPipe* and other methods using common cells and genes (Figure 3C). Overall, we found relatively high average Pearson correlations (nearly all above 0.9), with correlations between *scPipe* and *scruff* the highest, whereas, with *celseq2* and *zUMIs*, the correlations were slightly lower. *Celseq2*’s use of a different aligner (*Bowtie2*) which aligns to the transcriptome might account for the lower correlations, while for *zUMIs* the inclusion of intron reads in the gene counts is partially responsible, with the correlation increasing when the run in exon-only mode. After cell-level quality control (see Methods), in terms of the overlap in cells detected by different methods, the majority (266) were common across all methods, with a further 38 found by at least 3 out of the 5 workflows tested (Figure 3D). *kallisto bustools* detected 78 unique cells. This unexpected filtering result should on account of that cells with more detected features by *scPipe* had fewer features for *kallisto bustools* (Figure 3E). Next, we applied *GLMPCA* (38), an alternative dimension reduction method for visualizing the raw counts from scRNA-seq data, with example plots shown in Figure 3F. Clear separation between cells from the different cell lines was observed for all preprocessing workflows. Samples from cell lines H1975 and HCC827 were observed to be more similar according to their bulk expression profiles in a previous study (39), which is broadly consistent with what we observe here at the single-cell level.

### Gene quantification for 10x workflows

The same metrics were applied to the raw count matrix to compare the performance of 7 preprocessing workflows applicable to droplet-based 10x datasets (*scPipe, kallisto bustools, salmon alevin, Cell Ranger, dropSeqPipe* and *Optimus*) after applying *emptyDrops* or *Cell Ranger* v2 filtering (see Methods) (Figure S2).

In contrast to the CEL-Seq2 results, no systematic quantification bias was observed between workflows across datasets (Figure 4A-C show results for the 10xv2_3cl, 10xv3_5cl, and 10xv2_tissue2 datasets). The number of cells identified varied between preprocessing workflows, with *zUMIs* and *kallisto bustools* calling the highest number for the 10xv2_3cl dataset, *Optimus* for the 10xv3_5cl dataset and *salmon alevin* for the 10xv2_tissue1 and 10xv2_tissue2 datasets (Figure 4A).

**Fig. 4.**
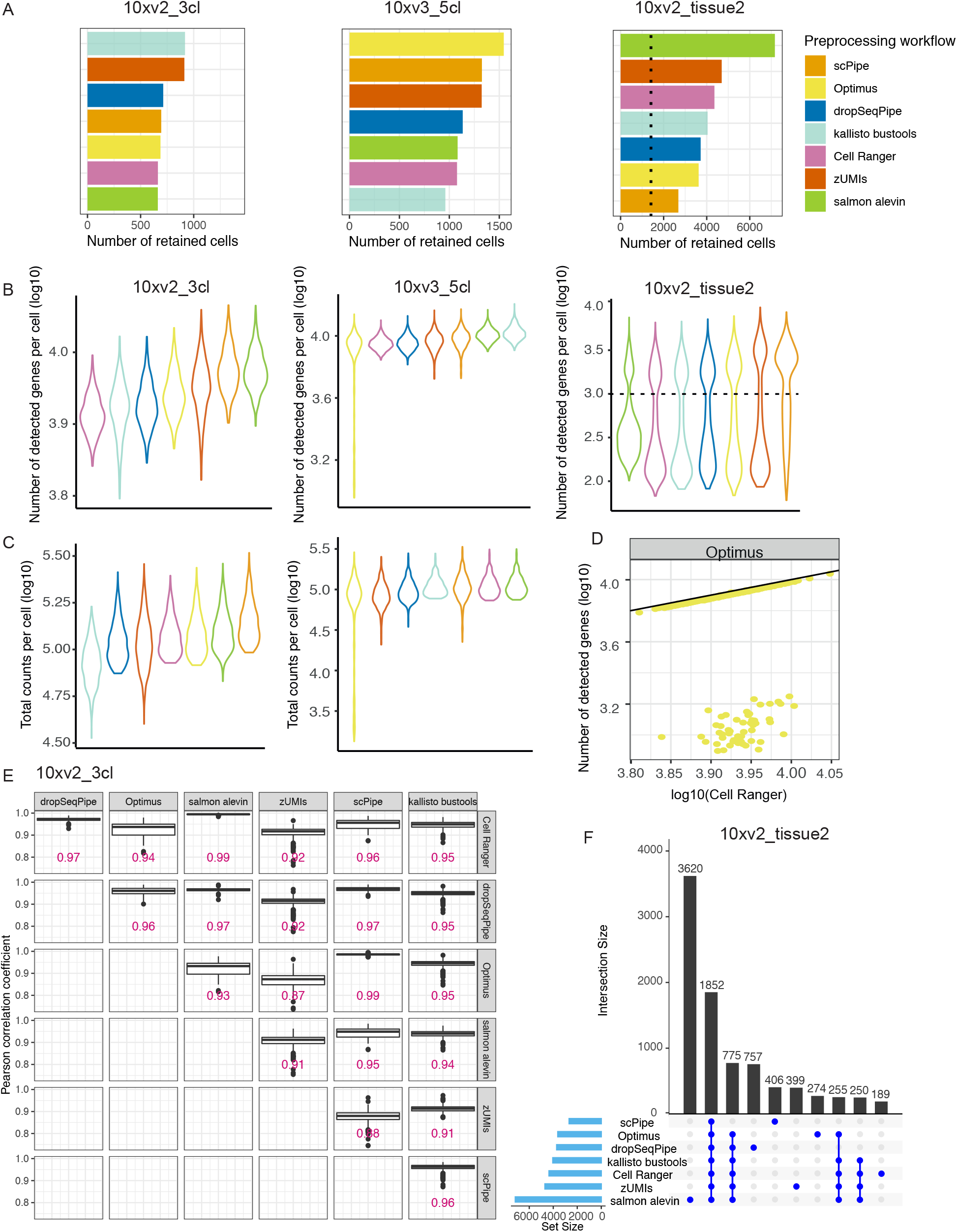
Comparing gene expression quantification of different scRNA-seq preprocessing workflows on droplet-based datasets. **A)** The number of cells detected are plotted on 10xv2_3cl, 10xv3_5cl and 10xv2_tissue2 datasets accordingly. The black dashed line represents the number of cells provided with a label. **B)** Number of detected genes per cell and **C)** total counts per cell (both on a log 10-scale) are shown. Here, since a bimodal distribution is displayed on the 10xv2_tissue2 dataset, to assist visualization, a dashed black line is drawn at detecting one thousand genes per cell. **D)** The number of detected genes per cell of *Optimus* are plotted against those from *Cell Ranger* on 10xv3_5cl dataset. And on the 10xv2_3cl dataset, the Pearson correlation coefficients of genes between each pair of preprocessing workflows are shown in **E)**. In terms of obtained cells after filtering across workflows, **F)** the UpSet plot is used to display intersections of retains cells across workflows on 10xv2_tissue2 dataset.

The number of detected genes and total counts per cell (Figure 4B-C) also varied by dataset. We compared these metrics obtained by *Cell Ranger* versus other workflows in a pair-wise manner (Figure S3A-D). For the 10xv2_3cl dataset, *zU-MIs* and *salmon alevin* detected more genes per cell. In terms of total counts per cell, *Cell Ranger* generally recovered higher counts in cells possessing lower total counts than other workflows (slope of the linear relationships above 1). *kallisto bustools* systematically underestimated the total counts per cell, which might be related to its naive UMI collapsing strategy. For the 10xv3_5cl dataset, the 7 methods are relatively concordant (Figure 4B-C), with the exception of *Optimus* which underestimates the number of genes and total counts in a subset of cells (Figure 4D), indicating that it may not be appropriate to use Levenshtein edit distance to correct CB from the extended 10xv3 barcode list. For the mouse tissue datasets, *zUMIs* detected more genes per cell and together with *scPipe*, recovered higher counts per cell, while other workflows were broadly similar. The *dropSeqPipe* software performed similarly to *Cell Ranger* across all datasets, which is to be expected as they apply similar methods at each step.

Calculation of the Pearson correlation of expression for common cells across workflows showed high average pairwise concordance for all workflows, especially between *salmon alevin, dropSeqPipe* and *Cell Ranger* (Figure 4E, S3E), with mean correlation consistently above 0.95. *scPipe* and *Optimus* also displayed high correlations with *dropSeqPipe* on single cell datasets. However, *zUMIs* showed relatively lower correlations with other workflows. Calculating correlations with intron counts excluded by *zUMIs* didn’t noticeably increase the correlation values, which indicates that the addition of intron counts is not a major cause for the lower correlations (Figure S3F).

Next, we filtered out low quality cells based on median absolute deviation using *scater* (see Methods). *UpSet* plots were generated to assess the concordance of retained CBs after filtering across workflows (Figure 4F, S3G). We found that most of the retained cells were common cells across workflows. An exception was for the 10x tissue datasets, in which *salmon alevin* retained 3.6k cells unique to this workflow, which was two-fold more than the common cells (1.8k) found between all workflows while all other workflows with the exception of *kallisto bustools* identified between 189 (*Cell Ranger*) and 775 (*dropSeqPipe*) unique barcodes that were not observed by other methods. These unexpected many unique cells should result from the step calling genuine cells. Cell calling method*emptyDrops*, which does not apply a simple threshold to distinguish real cells provides results rely on profiles of ‘cells’ with low UMI content. On tissue datasets, different preprocessing workflows might provide distinct information of droplets with low content, leading to varied sets of cells to be retained across preprocessing workflows.

### Comparing gene biotype detection across scRNA-seq preprocessing workflows

Genes of different biotypes can have systematically distinct length distributions and sequence similarity (40) and current RNA-seq tools have been shown to quantify genes possessing these characteristics differently (41, 42). To explore differences in the detection and quantification of genes biotypes across workflows, we investigated the signal and noise characteristics stratified by biotype.

#### Gene biotype detection for CEL-Seq2 workflows

For the representative plate_3cl dataset, the highest sensitivity and most lncRNAs were obtained with *celseq2*, which also detected features in the misc_RNA biotype (Figure 5A), while *zU-MIs* detected fewer pseudogenes compared to other methods. Density plots of the total counts per gene (Figure 5B) were similar for all methods, except for *kallisto bustools*. To compare the biological noise for each workflow, the biological coefficient of variation (BCV) was calculated using the known cell labels from each dataset as the ground truth (see Methods). BCV measures the proportion of gene expression attributable to biological variability (43), and typically starts from higher values at low abundance and decreases monotonically as gene abundance increases.

**Fig. 5.**
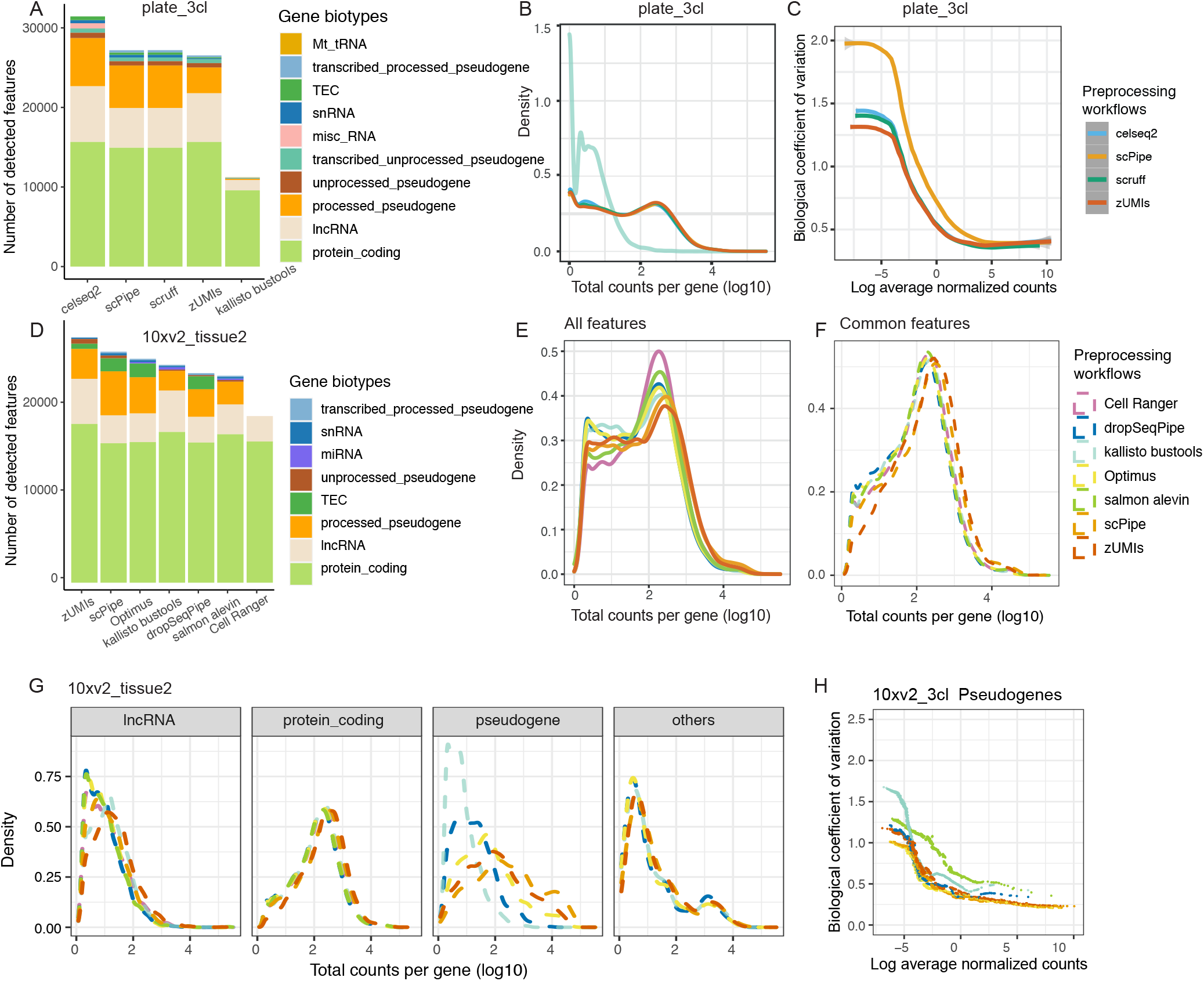
Comparing gene biotype detection of different scRNA-seq preprocessing workflows on droplet-based datasets. For plate-based datasets, **A)** number of detected features per gene biotype, and **B)** density of total counts per gene (on a log 10-scale) are shown on plate_3cl dataset. The biological coefficient of variation (BCV) for each feature was calculated and plotted against gene abundance in **C)** on plate_5cl1 dataset. For droplet-based datasets, **D)** number of detected features per gene biotype, and density of total counts per gene of **E)** all features and **F)** common features across workflows (on a log 10-scale) are shown on 10xv2_tissue2 dataset. Next, **G)** density plot were break down by features for gene biotypes lncRNA, protein coding genes and pseudogenes from common cells. The BCV for each feature of pseudogene was calculated and plotted against gene abundance in **H)** on 10xv2_3cl dataset. Colors represent different preprocessing workflows.

We observed such trends for most listed preprocessing workflows with selected biotypes using common cells and all features (Figure 5C), with the BCV trend for *scPipe* systematically higher than the trends for other methods, which are more consistent (results for *kallisto bustools* were excluded from this plot due to the limited numbers of features available to estimate BCV reliably).

#### Gene biotype detection for 10x workflows

Results for the 10xv2_tissue2 dataset (chosen as a representative example, Figure 5D-G) show that the number of detected genes and proportion of counts assigned varies across different biotypes between workflows, with some workflows more likely to detect or assign counts to a particular class of genomic features than others (Figure 5D, S4A). Most of the counts are delivered as protein coding genes across workflows. *Cell Ranger*’s use of a curated reference annotation restricts analysis to lncRNAs and protein coding genes, while *scPipe*, followed by *Optimus* assigned fewer counts to protein coding genes and together with *zUMIs* delivered a greater proportion of counts to pseudogenes. For short non-coding RNA, *scPipe, Optimus* and *zUMIs* also returned more miRNAs, while *kallisto bustools* and *salmon alevin* assigned a higher proportion of counts to the misc_RNAs on single cells and tissue cells datasets respectively. This influenced the count distributions as shown in Figure 5E and S4B, with lncRNAs, pseudogenes, and ‘other’ classes contributing more to the lower abundance peak and protein coding genes to the higher abundance peak. *Cell Ranger* detected fewer genes at the lower peak and more genes at the higher peak consistently, whereas other methods provided relatively fewer features in the higher peak (Figure 5E and S4B). On the 10xv2_3cl dataset, *kallisto bustools* displayed an obvious third intermediate peak (Figure S4B). If we restrict our analysis to features that are common across workflows, the lower peak is less prominent (Figure 5F), suggesting that the features quantified with low abundance tend to be discordant between workflows, which is in agreement with previous results (41).

Figure 5G shows how the different peaks are dominated by distinct biotypes, with protein coding genes displaying similar densities with a peak in the high abundance range, whereas genes of other biotypes are distributed more at the low abundance end of the distribution. Especially for common pseudogenes, *kallisto bustools* displayed the sharpest left-skewed peak, suggesting more lowly expressed pseudogenes yielding fewer than 10 counts were quantified with it. While for other workflows, the peak shifted from left to right, and a wider range was observed, in the order of *dropSeqPipe, Optimus, zUMIs* and *scPipe*, indicating the existence of workflow-specific quantification effects for pseudogenes.

We examined long-read transcriptome sequencing data on the same cell line mixture samples (44) to obtain an independent measure of pseudogene abundance in these data. Longer reads should map less ambiguously to pseudogenes compared to short-read data (45, 46). Comparing the proportions of counts mapped to pseudogenes from the different pre-processing methods to those obtained in the long-read data, which themselves varied between the 10x v2 and v3 chemistry, we consistently observed fewer counts being assigned by *dropSeqPipe* and the pseudo-alignment tools (*kallisto bustools* and *salmon alevin*), which suggests that these workflows systematically underestimate pseudogene abundance (Figure S4C). *Optimus* and *zUMIs* recover pseudogene count proportions that are similar to the long-read estimates, while *scPipe* systematically assigns more reads to pseudogenes and is probably overestimating signal in this class of features.

Given the differences in detecting specific biotypes across the 10x preprocessing workflows compared, we next investigate the biological information captured by the 3 most abundant classes, which are protein coding genes, lncRNAs, and pseudogenes respectively (Figure S4D). BCV and silhouette widths were calculated using the known cell labels from each dataset as the ground truth (see Methods) to compare the biological noise and signal across biotypes for each workflow, respectively.

We observed expected BCV trends (i.e. BCV decreases as abundance increases) for all listed preprocessing workflows for the selected biotypes using common cells and all features (Figure S5A-B). For single cell datasets, the BCV trends were highly similar for protein coding genes and lncRNAs (Figure S5A), although the BCV was systematically higher for the lncRNAs. The trends were also fairly consistent for the pseudogenes (Figure 5H), except for *salmon alevin*, where the BCV was systematically higher across the full range of abundance levels and *kallisto bustools* where the trend was markedly higher for the low abundance features only. For the tissue datasets, besides the higher BCV observed for pseudogenes with *salmon alevin*, there were differences across all biotypes between *scPipe, kallisto, zUMIs* and the rest on the workflows, where we observe higher variance of lowly expressed gene delivered by *scPipe, kallisto, zUMIs* (Figure S5B). Restricting the analysis to common features and cells detected across all workflows saw similar trends (Figure S5C-D), although BCV values decreased overall compared with the results from common cells (Figure S5A-B), suggesting that quantification of disconcordant features is a major source of variation between workflows.

Next, we investigated the biological signal recovered by features from specific biotypes. Silhouette widths calculated on GLMPCs, which measures how similar a cell is to its ‘known’ (pre-labeled) cell type compared to other cells were used and compared (see Methods). For both 10x single cell and tissue datasets, protein coding genes showed similar and higher silhouette widths, indicating biological signal was universally retained by all workflows (Figure S5E). The separation between different cell types can be visualized using t-SNE plots created using protein coding genes (Figure S6A-F, left-hand column). Most lncRNA and pseudogenes had silhouette widths above 0 for the single cell datasets (Figure S5E, left panel), which is indicative of a signal that can also be visualized in the t-SNE plots (Figure S6A-C, middle and right columns). However, on the tissue datasets, silhouette widths above 0 were only observed for lncRNAs with *zUMIs* and for pseudogenes, with *scPipe, Optimus* and *zUMIs* (Figure S5E, right panel) which can also be seen in the t-SNE plots (Figure S6D-E, middle and right columns), suggesting that the separation between the known cell groups was well less defined by these feature types. Although *kallisto* and *salmon* were recommended for detecting lncRNAs in a previous bench-marking study (47), we did not observe this preference consistently across our datasets.

Additionally, looking at t-SNE plots based on pseudogenes for *kallisto bustools* (Figure S6B) and *salmon alevin* (Figure S6E) shows a less clear separation between cell types compared to those provided by other workflows or for other biotypes, which suggests that pseudo-aligners quantified pseudogenes using 3’ sequencing data present less clear biological information.These results suggest that focusing analysis efforts on the signal from protein coding genes in datasets that profile complex tissues with greater cell type diversity may be an optimal strategy.

### Comparing the performance of different combinations of preprocessing workflows and normalization methods

We next examined the degree to which the choice of processing workflow influences downstream analysis. Specifically, we look into how preprocessing impacts normalization, highly variable gene (HVG) selection and clustering.

#### Normalization methods and evaluation metrics

Normalization has been shown to be an influential step in previous benchmarking studies (27, 48). We applied five popular and well proven normalization methods (5, 49), including *DESeq2* (50), *scone* (49), *scran* (51), *Linnorm* (52) and *sctrans-form* (53) to explore how well different approaches remove any of the inherent biases introduced by preprocessing.

Silhouette widths of known cell groups and unwanted variation explained by library size and wanted variation explained by known cell groups on principal components (PCs) after normalization was used to evaluate this. To summarise the results across the many different combinations of dataset × pre-processing method × normalization algorithm, linear models were fitted with silhouette widths as the response variable and the different methods as covariates. Higher silhouette widths, and reduced unwanted variation represents better performance (see Methods). PCA plots were also generated for each combination of preprocessing workflow and normalization to assist in visualizing the data structure. Moreover, after normalization, HVGs were selected for each combination of dataset × preprocessing method × normalization algorithm using *scran* (see Methods).

#### Normalization performance assessment on CEL-Seq2 datasets

Considering silhouette widths on the CEL-Seq2 RNA mixture datasets, no preprocessing workflow systematically outperformed others when combined with different normalization methods (Figure 6A left column). Evaluation of inspecting PCA plots agrees with silhouette widths and example PCA plots are shown in Figure S7A. Most of the combinations presented expected trajectory paths, and combinations evaluated with higher silhouette widths displayed better separation between distinct RNA mixtures, e.g. *scruff*, followed by *scPipe* combined with *scone*. In terms of variation explained by library sizes, relatively less variation was explained by *scPipe* combined with any normalization methods compared to other preprocessing methods, which is preferred (Figure S7B).

**Fig. 6.**
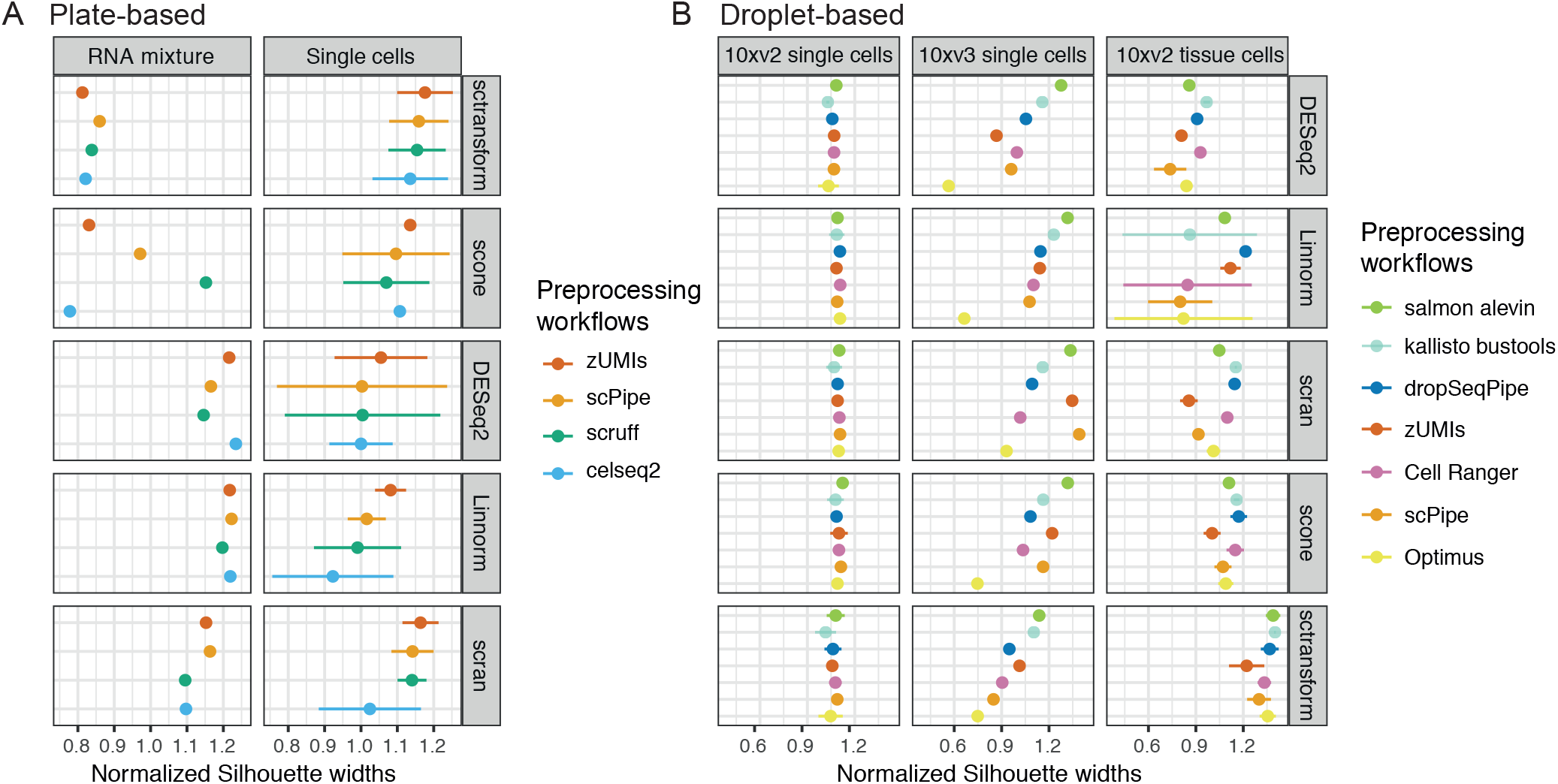
Comparing performance of combinations of different scRNA-seq preprocessing workflows and normalization methods. Silhouette widths are calculated based on known cell labels after applying different normalization methods and normalized against the silhouette widths obtained without any normalization. Dot plots (mean silhouette widths ± s.d) are shown on the plate-based datasets in **A)** and on the droplet-based datasets in **B)**. Common cells between workflows were used in the tissue datasets. Colors denote different preprocessing workflows.

For the single cell datasets, in terms of silhouette widths, *zU-MIs* performed relatively better, while *celseq2* was slightly worse on average across all normalization methods (Figure 6A right column and S7C). Regarding unwanted variation, performance was fairly similar across combinations (Figure S7D).

#### Normalization performance assessment on 10x datasets

Highly similar silhouette widths after normalization were observed on the 10xv2 single cell datasets (Figure 6B, left-most column). However, for the 10xv3_5cl dataset, silhouette widths varied considerably between preprocessing workflows after normalization (Figure 6B, middle column). On the 10xv3_5cl dataset, *salmon alevin* produced consistently high silhouette widths across normalization methods, followed by *kallisto bustools*, while *Optimus* consistently gave the lowest values (Figure 6B, middle column), which is likely caused by the under-quantification of gene expression in a subset of cells. Considering unwanted variance, in accordance with evaluation based on silhouette widths, *Optimus* retained the more variance by library size, while other methods performed fairly similarly (Figure S8A).

Inspection of PCA plots (Figure S7B) from scran-normalized data showed that combinations with higher silhouette widths (*scPipe, zUMIs, salmon alevin*) had better separation of H1975 and HCC827 cells compared to other workflows, although all methods showed good separation overall.

For 10xv2 tissue datasets, the number of cells retained by different preprocessing methods varied considerably (1-to 7-fold variation beyond the number of cell type labels provided, Figure 4A right-most column) so silhouette widths were calculated for all cells (Figure S8C) and common cells between workflows (Figure 6B, right-most column). Analysis of common cells gave higher silhouette widths that were less variable compared to the silhouette widths calculated using all cells, indicating that the unique cells recovered by different workflows added considerable noise to the analysis (Figure S8C). Overall, *zUMIs* and *scPipe* tended to have slightly lower silhouette widths, while the other workflows performed fairly similarly (Figure 6B right-most panel, S8D). Taking median explained variance into account, *scPipe* and *zUMIs* were also not preferred because they retained more unwanted library size variation (Figure S8E). The two pre-processing workflows that applied pseudo-alignment tools, *salmon alevin* and *kallisto bustools* in contrast retained less unwanted variation.

Additionally, considering the rank of these preprocessing workflows across different normalization methods on both single cell and tissue datasets based on silhouette widths (Figure 6B), we found that there was no consistent winner. However, the ranks were relatively stable on average, e.g., on tissue datasets, *dropSeqPipe* and *kallisto bustools* are always better and *scPipe* and *zUMIs* are slightly worse, suggesting that individual preprocessing workflows might incorporate quantification biases that cannot be eliminated by normalization.

#### Summary of normalization results

The performance evaluated by silhouette widths is summarized in Figure S9. Although a wide range of silhouette widths was observed among different workflows combined with different normalization methods, all displayed increased silhouette width compared to the results obtained by different preprocessing methods without normalization, suggesting that all normalization methods are highly effective on these data. In terms of preprocessing workflows, although very similar performance is displayed across methods, normalization methods perform slightly better with *salmon alevin* on droplet-based datasets, with relatively less variation between normalization algorithms. Following closely behind is *dropSeqPipe, kallisto bustools*, then *zUMIs, Cell Ranger, scPipe* and finally *Optimus* which shows the most variation in performance. On plate-based datasets, less variation is shown with *zUMIs*, followed closely behind by *scruff, scPipe* and *celseq2*. Looking at the results from the normalization side, *sctransform* combined with any preprocessing workflow delivers better results across datasets on average, which corroborates with earlier findings (48, 54), followed closely behind by *scran* and *scone* whereas, *DESeq2* and *Linnorm* performed slightly worse than the other methods with more variation in performance.

#### Performance assessment with HVGs on 10x datasets

We then selected HVGs (see Methods) for each combination of dataset × preprocessing method × normalization algorithm. Although the proportion of HVGs of different biotypes varies widely across datasets, protein coding genes account for the largest proportion of HVGs (Figure S10A-B). Proportions also vary among preprocessing workflows. *Cell Ranger* delivered more protein coding genes than other workflows as expected since it excludes other biotypes, while *scPipe* followed by *Optimus* returned more pseudogenes (Figure S10A-B). While most of the protein coding HVGs are detected in common between workflows, there are lists of unique genes (more than 100 genes) retained by individual preprocessing workflows on both the single cell (Figure S10C) and tissue cell datasets (Figure S10D), e.g. *salmon alevin, zUMIs* normalized by *scran* on single cell datasets. For the lncRNAs and pseudogenes biotypes, the overlap of HVGs between all methods was modest by comparison with 21 and 4 features respectively for the single cell data (Figure S10C).

On the tissue datasets, nearly every preprocessing workflow yielded a unique list of HVGs. For example, after normalization by *scran*, the number of genes in the unique list of *scPipe* was 411 for protein coding genes and 158 for pseudogenes (Figure S10D). For lncRNAs, only 5 HVG features were in common between all workflows, and for the pseudogenes, there were no common highly variable features (Figure S10D). The ability of these HVGs to recover the expected data structure after individual preprocessing and normalization combinations was once again evaluated by silhouette widths (Figure S11). For the simpler single cell datasets, little difference was observed between methods, while on the tissue datasets, the performance improved after feature selection, with the variation between workflows decreasing, indicating that feature selection helps remove noise and variance that obscures data structure across workflows.

### Comparing the performance of different combinations of preprocessing workflows and clustering methods

#### Clustering methods and evaluation metrics

There has been much research focused on the performance of clustering methods in terms of sensitivity of parameters, accuracy, robustness, etc. (48, 55, 56). Here, we aim to investigate the impact of preprocessing on clustering instead of ranking clustering methods based on their performance. We selected representative clustering methods implemented in R and evaluated their performance when they reach the expected number of clusters based on the labels available. *RaceID* (57), *SC3* (58), *scran* and *Seurat* (59) were methods included. Both classic unsupervised methods and combined Support Vector Machine (SVM) methods in *SC3* were used. Graph-based clustering methods have been shown to perform fairly well previously, so *scran* with algorithms of fast-greedy, louvain and walktrap and *Seurat* with louvain and SLM (60) were all included in the evaluation. Here, *Seurat* is applied using its default pipeline according to the recommended vignette after the normalization step (that includes normalization). The entropy of cluster accuracy (ECA), the entropy of purity (ECP) (see Methods) and adjusted Rand index (ARI) (61) were used for assessment considering intra-cluster similarity, external criterion, and similarity of clustering partition with known clusters, respectively. ANOVA was then applied to assess the relative variation explained by the main analysis steps (preprocessing, normalization and clustering, see Methods). We also fitted a linear model to evaluate the extent to which specific methods or workflows at each analysis step, including preprocessing, normalization, and clustering, influenced the clustering result. ARI, ECA, or ECP were used as dependent variables in this analysis. Considering the intrinsic difference between experimental designs, ANOVA and linear model were fitted to the various combinations of methods separately for datasets with different designs.

#### Clustering performance assessment on CEL-Seq2 datasets

The expected clustering results for the RNA mixture datasets are different from that of the other datasets due to the inbuilt trajectory paths and labels provided via the mixture proportions. In terms of ARI, combined with selected normalization and clustering methods, *scruff* produced more combinations with better performance, while *scPipe* provided relatively fewer (Figure 7A). However, *scran* clustering using walk-trap and louvain performed best overall when combined with *Linnorm* and either *scruff* or *scPipe* (Figure S12A). These combinations also performed best in terms of entropy (Figure S12B). The *Seurat* pipeline ranked closely behind and performed consistently across all preprocessing workflows (Figure S12A). Overall, *scruff* and *celseq2* were ranked first and second across all normalization and clustering method combinations according to the coefficients from the linear modeling, indicating better performance on average (Figure 7C). Inspection of t-SNE plots shows that combinations yielding better performance display clearer separation of cells from different mixture groups, whereas method combinations that perform relatively worse have clusters made up of cells from multiple mixture groups (Figure S12C).

**Fig. 7.**
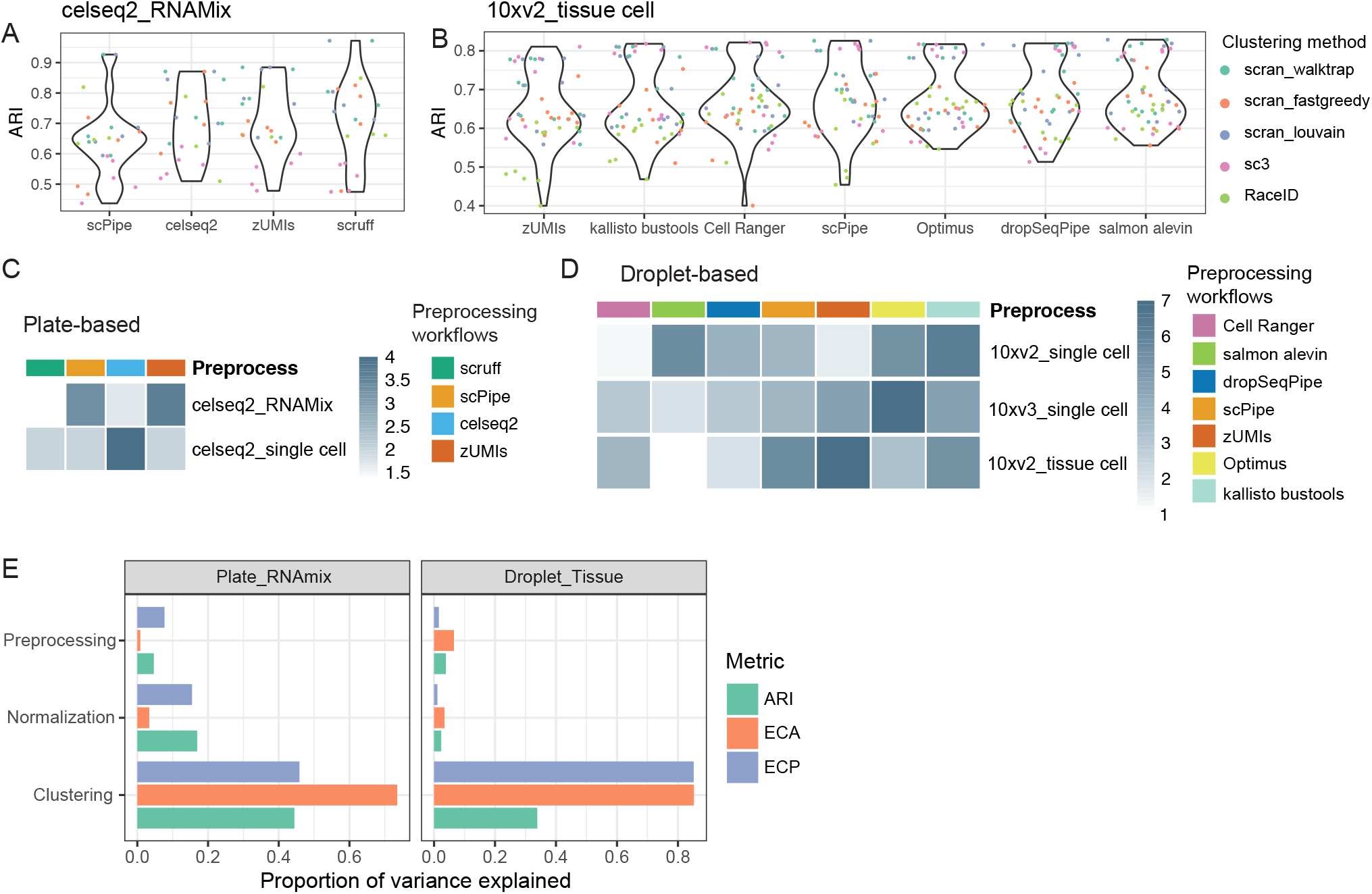
Comparing performance of combinations of different preprocessing, normalization and clustering methods. Performance evaluated by ARI are displayed in violin plots broken down by different preprocessing workflows on **A)** plate-based RNAmix and **B)** droplet-based tissue datasets (based on common cells). Each dot represent a combination, and is colored by the clustering method applied. Next, preprocessing workflows’ influence on clustering results is summarized. Results for plate-based data are in **C)**, and droplet-based data are in **D)**. Colors represent the rank of their weighted average rank across evaluation metrics (ARI, ECA, ECP). Lighter color means better performance (i.e. higher rank). ANOVA model based upon a given performance metric ~ preprocessing+normalization+clustering+experimental design was used to calculate the proportion of variance explained by main analysis steps. Results for the plate-based RNAmix data and droplet-based tissue data (using common cells) with ARI, ECA and ECP used as respective metrics are plotted in **E)**. Colors denote different evaluation metrics.

For the single cell datasets, clustering is a much easier task since these comprise relatively fewer (3 or 5) well-separated clusters. On these data, nearly all method combinations obtained ARIs of 1 (Figure S13A) and both ECA and ECP of 0 (Figure S13B), except for *SC3* with SVM. It is worth noting that *SC3* with SVM is suggested for use on datasets containing more than 5k cells, which does not suit our smaller plate-based datasets, so we removed combinations containing *SC3*-SVM before fitting any models to further summarise the results.

T-SNE plots of clustering results returned by different methods for the 5 cell line mixture dataset agree with the known cell types present (Figure S13C). Overall, *scruff, celseq2* and *scPipe* were ranked equal first and across all normalization and clustering method combinations using the linear model coefficients (Figure 7C). Considering the proportion of variance explained, we observed that for both the RNA mixture (Figure 7E, left-hand panel) and single cells datasets (Figure S13D), clustering methods had a greater influence on performance, followed by normalization methods, while pre-processing workflows explained the least variation in performance.

#### Clustering performance assessment on 10x datasets

The 10x single cell control datasets also have a simple structure (3 or 5 expected clusters), and the performance of most method combinations, evaluated using ARI values, reached 1 (Figure S14A). Correct clustering of cells can be observed via t-SNE plots (Figure S14B). We added up the number of combinations with both ECP and ECA at 0, and found that *Cell Ranger* and *salmon alevin* delivered more optimal combinations, followed by *scPipe*, indicating that these methods consistently delivered reliable results (Figure S14C). On average, *Cell Ranger* and *dropSeqPipe* rank amongst the top 3 in terms of performance as estimated by the coefficients from the linear modeling (Figure 7D).

For the tissue datasets, clustering results generated based on common cells display similar performance, with comparable ARIs (Figure 7B and S15A). Overall, *salmon alevin* and *dropSeqPipe* rank amongst the top 2 for performance as assessed by the coefficients from the linear model analysis (Figure 7D and Figure S15B). Consistent with observations from the plate-based datasets, the proportion of variation in performance explained in the tissue dataset was greatest for clustering methods (Figure 7E, right-hand panel). The variation in performance explained by normalization and preprocessing methods were both negligible.

## Discussion

### Summary of performance of preprocessing workflows

We compared the performance of 9 preprocessing workflows across CEL-Seq2 and 10x Chromium platforms on datasets with varying biological complexity, and explored their quantification characteristics and impact on downstream analysis. In terms of preprocessing workflows designed for CEL-Seq2, the workflows compared showed high concordance in quantification, with small discrepancies on detected features. Among them, *celseq2* is more sensitive and returned more non-protein coding genes. Very similar results were returned by *scPipe* and *scruff* which share a lot of the same preprocessing choices, although more biological noise was onserved in lowly expressed genes by *scPipe* due to its UMI quantification approach. *scPipe* only collapses UMIs that differ by a hamming distance of 1 with more than 2-fold difference in counts, which means the number of UMI counts delivered by *scPipe* is relatively higher. In terms of downstream analysis, on the simpler datasets (single cell mixtures), nearly all workflows produced clustering results that agreed well with the known cell type labels combined with any normalization and clustering methods. On the more complex RNA mixture dataset, with clusters that are less well separated due to the gradient of mixture proportions used in this design, *scruff* performed best, followed by *celseq2* and *scPipe* (Figure 7C). Our analysis revealed that *kallisto bustools* (v0.39.3) was unsuitable for use on CEL-Seq2 data due to its UMI handling strategy, which limits the dynamic range that can be observed in the presence of short UMIs (6bp in the case of CEL-Seq2). This approach does not cause any noticeable feature detection issues in protocols which use longer UMIs such as 10x Chromium (10-12bp).

As for the workflows designed for the 10x Chromium platform, regarding detection and quantification of genes, several differences were observed. *Cell Ranger* provided more features with higher abundance, which are mostly made up of protein coding genes, whereas other workflows displayed a second peak of features with lower abundance. *kallisto bustools* delivered more features at an intermediate abundance level on the 10xv3_3cl dataset.

Also, we found that *kallisto bustools, scPipe* and *zUMIs*) retained more biological noise, particularly amongst lowly expressed features compared to other workflows analyzed on the tissue datasets. Many of the lowly expressed features were not found in common between preprocessing workflows. These differences indicate the uncertainty in detecting lowly expressed features.

Previous studies have illustrated *Cell Ranger*’s bias for genes with low uniqueness (24) and the adverse effect of running it with a full annotation file (30). *Cell Ranger*’s use of a targeted reference annotation that focuses on protein coding genes and lncRNAs and excludes biotypes that are more difficult to resolve with short-read sequencing, such as small RNAs and pseudogenes, would likely benefit other preprocessing workflows. Use of a modified feature set would not only alleviate the impact caused by multi-mapping, but also avoid the workflow-specific gene quantification impacts seen in other gene biotypes that are not of principal interest (such as pseudogenes). This would obviously not suit studies in which quantification of such features are of particular interest. We observed that biological signal present in the raw counts were nearly identical between *Cell Ranger* and the other preprocessing workflows compared. Additionally, after downstream analysis, we observed *Cell Ranger*’s performance to be consistently high and relatively stable, which supports the results of another recent study (30). Some pre-processing workflows were found to be relatively more or less likely to assign counts to a particular class of genomic features than others. For example, *scPipe* assigned systematically more reads to pseudogenes, whereas *dropSeqPipe, salmon alevin* and *kallisto bustools* assigned many fewer reads to this feature class. In addition, for the two workflows that use pseudo-alignment, *kallisto bustools* provided more pseudogenes yielding a limited number of counts, and *salmon alevin* retained more biological noise in pseudogenes.

After normalization, evaluating the performance of preprocessing workflows based on known biological information uncovered similar performance in all but a few method combinations, indicating high concordance between preprocessing workflows. After selecting highly variable genes, discrepancies between workflows was further reduced. Interestingly, nearly every workflow revealed a unique list of features that included varying proportions of different gene biotypes.

Regarding clustering results, similar performance was observed across different workflows and methods on the single cell datasets. On tissue datasets, where experimental complexity was higher, we still observed similar performance when clustering analysis was restricted to common cells identified across all workflows. *Cell Ranger, dropSeqPipe* and *salmon alevin* performed slightly better on droplet-based datasets on average. Although slightly lower correlations were observed between *zUMIs* with other workflows, and it uniquely considers the intron counts within selected workflows, we did not observe improved clustering results by applying it. Intron reads are indicated to be informative as they likely originate from nascent mRNA (62) and were showed to assist in extracting more information when included in quantification (17). However, the scRNA-seq data we analysed both rely on poly(A) selection, which may limit the amount of intron signal that can be extracted.

#### Limitations

Our benchmarking study is subject to several limitations. First, although there are other protocols such as InDrops, Drop-seq, Sort-seq etc., and different protocols are known to influence downstream analysis (26, 54), our results are restricted to datasets from two protocols, CEL-Seq2 and 10x Chromium. Second, we did not expand our analysis to to include other important aspects of the single-cell data processing, such as dimensionality reduction, feature selection, etc. (48). Extending our benchmarking to cover additional protocols, new preprocessing methods and other data analysis tasks is left as future work.

Additionally, for QC we applied the same cell identification and cell-level filtering method based on soft thresholds in *scater*. This resulted in more retained cells in the tissue datasets, e.g., on the 10xv2_tissue2 dataset, only about 1.5k cells were provided with a cell type annotation; however, with our cell identification and filtering strategy, most preprocessing workflows recovered nearly 4k cells. The additional cells were found to contain more noise based on our evaluation metrics.

Furthermore, the available ground truth in the simple mixture experimental designs, which involved either mixtures of RNA or single cells from 3 to 5 cell lines, are less complex than can be expected in regular experiments. Only the tissue cell datasets are representative of the biological complexity found in research samples. The sample types studied are limited to cells from cancer cell lines or primary lung tissue. Further work could explore the performance of pre-processing workflows on datasets that include a more diverse range of cell types, e.g., immune cells or cells that make up the tumor microenvironment, although we would anticipate broadly similar results.

Another aspect not investigated in our study is which specific step within different preprocessing workflows has the most influence on performance. Although we observed quantification differences between workflows, we did not delve further into the individual steps within a workflow, which include alignment, deduplication, etc. Previous studies have already provided relatively comprehensive comparisons of alignment and quantification methods (27, 63). Hence, the results from our benchmarking study are targeted more to workflow users rather than workflow developers.

#### Summary of Findings

Our assessment investigated the quantification performance of preprocessing workflows and their impact on downstream analysis. We found that scRNA-seq preprocessing workflows varied in their detection and quantification of lowly expressed genes across datasets. However, after subsequent downstream analysis by well-performing normalization and clustering methods, even if the proportion of different gene biotypes detected differed and workflow-specific genes were identified within the various sets of highly variable genes, nearly all combinations delivered good performance with relatively minor differences in the final cell clustering results. Our detailed analysis of more than 2,000 dataset × method combinations, made possible by the *CellBench* evaluation framework, finds that the choice of preprocessing workflow has relatively less impact on the results of a single-cell analysis than subsequent downstream analysis steps such as normalization and clustering.

## Methods

### Preprocessing workflows compared

We evaluated 9 publicly available preprocessing workflows in total (Table S1). Workflows that started from raw `FASTQ` files and pro-vide a cell-by-gene count matrix as output were chosen. All workflows were installed and run locally, except for *Optimus*, which was run on Terra (https://app.terra.bio).

For all analyses, the genome, transcriptome (both cDNA and ncRNA) and `GTF` versions used for human datasets was Ensembl GRCh38, release 98 and for mouse datasets Ensembl GRCm38, release 99. The software versions used were as follows: *Cell Ranger* (v3.0.2), *celseq2* (v0.5.3.3), *dropSeqPipe* (v0.4.1) (YouTube tutorial link https://www.youtube.com/watch?v=4bt-azBO-18), *kallisto* (v0.46.0), *bustools* (v0.39.3), *Optimus* (v1.3.6 for 10x v2 datasets and v2.0.0 for 10x v3 datasets on Terra), *salmon* (v1.0.0), *scPipe* (v1.8.0), *scruff* (v1.4.2), *zUMIs* (v2.5.5), *bowtie2* (v2.3.4.1), *Samtools* (v1.9), *STAR* (v2.6.1c) and *Rsubread* (v2.0.0). Parameters within each preprocessing workflow were selected as recommended in the user guides. More details of each preprocessing workflow for each dataset is available at https://github.com/YOU-k/preprocess.

To compare computational performance, we created subsets of the data of varying sizes and carried out each analysis on a high-performance cluster (Intel(R) Xeon(R) CPU E5-2690 v4 @ 2.60GHz). We required 1 node and 8 PPNs for each run and claimed 8 cores in the scripts if there were parameters that allowed this; if not, 8 threads were claimed. We ran each workflow three times on each dataset and calculated the mean and standard deviation of the performance measures, which include CPU utilization, memory usage, and run time.

### Datasets

We selected 9 published datasets and created 1 new dataset for benchmarking (Table S2). The *scmixology* datasets are available from GEO under accession number GSE118767 (5). We selected plate-based CEL-Seq2 and 10x v2 droplet-based datasets containing cells from human lung adenocarcinoma cell lines that involved ‘pseudo-cells’ created by mixing cells (3 datasets) or bulk RNA mixtures (1 dataset) in different combinations using CEL-seq2 or actual single cells (1 dataset with 3 cell lines and 1 dataset with 5 cell lines). Annotation files for CBs with cell types were available at https://github.com/LuyiTian/sc_mixology.

The new dataset was created using single cells from the same five human lung adenocarcinoma cell lines (HCC827, H1975, A549, H838 and H2228) that were cultured separately. Cells were counted using Chamber Slides, and roughly 2 million cells from each cell line were mixed and processed by the 10x Chromium single cell platform using v3 chemistry. Afterward, libraries were sequenced on an Illumina Nextseq 500. Raw data from this experiment are available from GEO under accession number GSE154870. To generate its annotation files, *demuxlet* (https://github.com/statgen/demuxlet) was used to deconvolve cell identity via genetic information using the intermediate `bam` file obtained from the *scPipe* workflow.

The final dataset profiled mouse lung tissue from the Tabula Muris study, available under GEO under accession number GSE109774 (37). Raw `bam` files from channel `10X_P7_8` and `10X_P7_9` were downloaded and converted to raw `FASTQ` files by *bamtofastq* (v1.2.0) (https://support.10xgenomics.com/docs/bamtofastq). Annotation files for CBs with cell types were available from http://tabula-muris.ds.czbiohub.org.

The long-read Nanopore single-cell datasets used the same five human lung adenocarcinoma cell line mixture samples processed by the 10x Chromium single cell runs using v2 and v3 chemistry according to the protocols described in Tian *et al*. (44). The long-read based count-matrices provided by the authors were used to compare the abundance of particular gene biotypes with the matching short-read data.

### Cell quality control and cell annotation

In an attempt to standardize cell filtering, we applied *isOutlier* from the *scater* package (v1.14.6) setting library size, number of detected features, and percentage of mitochondrial genes per cell as filtering indicators with `nmads = 3` for all cell-by-gene count matrices created by different preprocessing workflows. Additionally, we applied `emptyDrops` from the *DropletUtils* package (v1.6.1) (14) (as recommended in (64)) to all selected 10x datasets. For the 10xv2_3cl dataset, *emptyDrops* failed to generate a reasonable number of cells when run on the *zUMIs* and *kallisto bustools* count matrices (i.e. around 60 thousand cells in each, which is far above expectation) so we used *Cell Ranger* v2 filtering to distinguish true cells for these 2 methods. The number of detected genes per gene biotypes, total counts per gene, total counts per cell, and Pearson’s correlation were all calculated based on the cell-by-gene count matrices obtained before filtering by *scater*. Here, the correlation was calculated using common cells and common features detected across all preprocessing workflows. GLMPCs were calculated using the *GLMPCA* package (v0.2.0) and *UpSetR* (v1.4.0) plots of cells found in common between preprocessing methods after filtering with using *scater* were generated for different datasets.

Because CB annotation files were generated based on one specific workflow, and the original analyses adopted different filtering strategies, it was possible to recover cells regarded as good quality by some preprocessing workflows that were not listed in the CB annotation files. Such cells were retained for further analysis and only removed upon calculating tailored evaluation metrics that required cell labels. Doublets detected by *demuxlet* in both the CEL-Seq2 and 10x single cell datasets were removed before normalization.

### BCV plot and biological signal on raw counts

For biological noise and signals, genes with specific biotypes were firstly extracted. Next, the biological coefficient of variation (BCV) was calculated using the filtered raw count matrix using `edgeR::estimateDispersion()` (v3.28.1) (65) and `trended.dispersion` was plotted using a `loess` line or directly with points. In the BCV plots, the x-axis displays the log-transformed counts obtained after *scran* normalization. Silhouette widths were calculated using the first 2 GLMPCs for single cell datasets and the first 20 GLMPCs for tissue datasets. To generate t_SNE plots, the same sets of genes across biotypes were extracted after normalized by *scran*. PCA was performed on them separately. The First 2 PCs on single cell datasets and the first 20 PCs on tissue datasets were used to create a t_SNE visualization.

### Data normalization

Five normalization methods were used to explore the impact the choice of preprocessing workflow has on this step and subsequent downstream analysis. The baseline *no normalization* option refers to the analysis of the raw counts directly without any further processing (this option was not used in downstream analysis). *DESeq2* (v1.26.0), *Linnorm* (v2.10.0) and *scran* (v1.14.6) were used with default settings. For *scone*, we set the maximum number of `RUVg` factors and maximum number of quality PCs both as 0. For *sctransform* (v0.2.1), we specified `1500` features as the number of variable features after ranking by residual variance.

To evaluate, we performed PCA with normalized counts firstly with default settings in *scater*. Next, silhouette widths were calculated with the first 2 PCs according to known cell clusters (provided by cell line identity or mixture proportion information) for all datasets except the tissue datasets. Considering the biological complexity of tissue datasets, silhouette widths were calculated using the first 20 PCs. Higher silhouette widths indicated better preservation of biological signals. Additionally, variance explained by library sizes and known cell groups on the first five PCs were summed up respectively as unwanted variation and wanted variation to assess whether known biological variation was preserved and confounded technical effects were well handled on all plate-based datasets and droplet-based single cell datasets. For the tissue datasets, variance explained from the first 20 PCs were partitioned into wanted and unwanted variation and then summed up.

### Clustering

Results from normalization were not directly applied with clustering methods. Except for *sctransform*, normalized counts were already selected with top 1.5k highly variable genes (HVGs), top 1.5k HVGs were obtained with `scran::modelGeneVar` and `scran::getTopHVGs`. Clustering methods from mainly four packages, *SC3* (v1.14.0), *Seurat* (v3.1.3), *RaceID* (v0.1.7), *scran* with *igraph* (v1.2.5) were used. To make it easier to interpret, we provided the number of clusters or specified related parameters with a range of values to reach the true value of the number of clusters. Other parameters were specified based on either the default settings or the author’s guidance from the user manual.

For *SC3*, both the classic unsupervised method and combined Support Vector Machine (SVM) method were used. *RaceID* parameters as suggested in the user reference manual were chosen. The required number of clusters were directly provided to *SC3* and *RaceID*. For *scran*, the number of nearest neighbors was specified at `5`, `10`, `30`, `50` and `100` to build a shared nearest-neighbour graph. Algorithms `fast greedy`, `Louvain` and `walktrap` in *igraph* were applied afterwards. For *Seurat*, we started from scratch using its own normalization method and clustered specifying resolution at `0.2`, `0.4`, `0.6`, `0.8`, `1.0`, `1.2`, `1.4` with the algorithms of `Louvain` and `SLM`. With *RaceID* and *SC3*, true number of cluster is provide. When applying *scran* and *Seurat*, only the result that produces a partition with the right or closest number of clusters was kept.

To evaluate clustering performance, the entropy of cluster accuracy (ECA), entropy of purity (ECP) (5) and adjusted Rand index (ARI) was used to assess intra-cluster similarity, external criterion, and similarity with known clusters, respectively. ECA measures the diversity of known cell groups within each cluster provided by given methods, and ECP measures the diversity of the cluster labels within each of the known true groups. Low values of ECA and ECP are favorable.

To summarize the results of each analysis, we performed ANOVA with the following model: metric ~ preprocess_workflow + norm_method + cluster_method + design. We also fitted a linear model using the `lm` function with the listed metrics as the dependent variable and the experimental designs and specific methods as covariates. The coefficient obtained for each method indicated to what extent these methods were influencing the clustering performance. Then, the average weighted rank of coefficients on preprocessing workflows across three metrics (ARI, ECA, and ECP) are calculated as a summary. Example t-SNE plots (created using *scater*) allowed visual assessment of different combinations’ performance. Other figures were created using *ggplot2* (v3.3.0) and heatmaps were created using *pheatmap* (v1.0.12).

### Benchmarking pipelines

*CellBench* (v1.2.0) was used to compare different methods as modules. The preprocessing workflows were individually applied to each dataset and the resulting cell-by-gene count matrix were input to `CellBench::apply_methods()` (36). Example code containing wrappers for use in *CellBench* for the methods compared are available at https://github.com/YOU-k/preprocess.

## ACKNOWLEDGEMENTS

This work was supported by funding from the National Health and Medical Research Council (NHMRC) Project Grant (No. GNT1143163 to M.E.R.), Fellowship No. GNT1104924 to M.E.R., the Chan Zuckerberg Initiative DAF, an advised fund of Silicon Valley Community Foundation (Grant Nos. 2018-182819 and 2019-002443 to M.E.R.), the Genomics Innovation Hub, Victorian State Government Operational Infrastructure Support, Australian Government NHMRC IRIISS and support from the Australian Cancer Research Foundation.

## Notes

### Competing Interest Statement

The authors have declared no competing interest.

