## Supplemental figures for "Benchmarking UMI-based single cell RNA-sequencing preprocessing workflows"

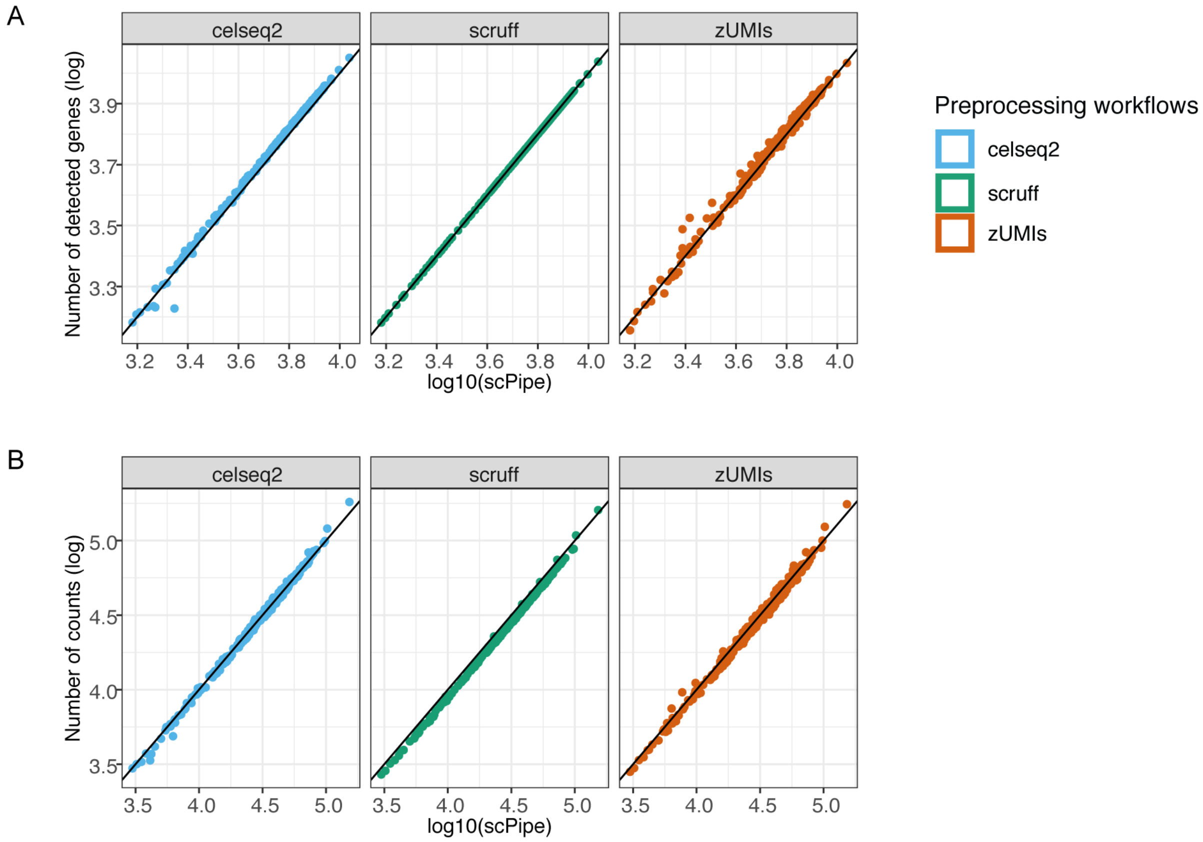


Figure S1, Comparing gene expression quantification of different scRNA-seq preprocessing workflows on plate_3cl dataset. In terms of common cells obtained across workflows, A) the number of detected genes per cell and B) total counts per cell of different preprocessing workflows are plotted against those from *scPipe* (both in a log10-scale).

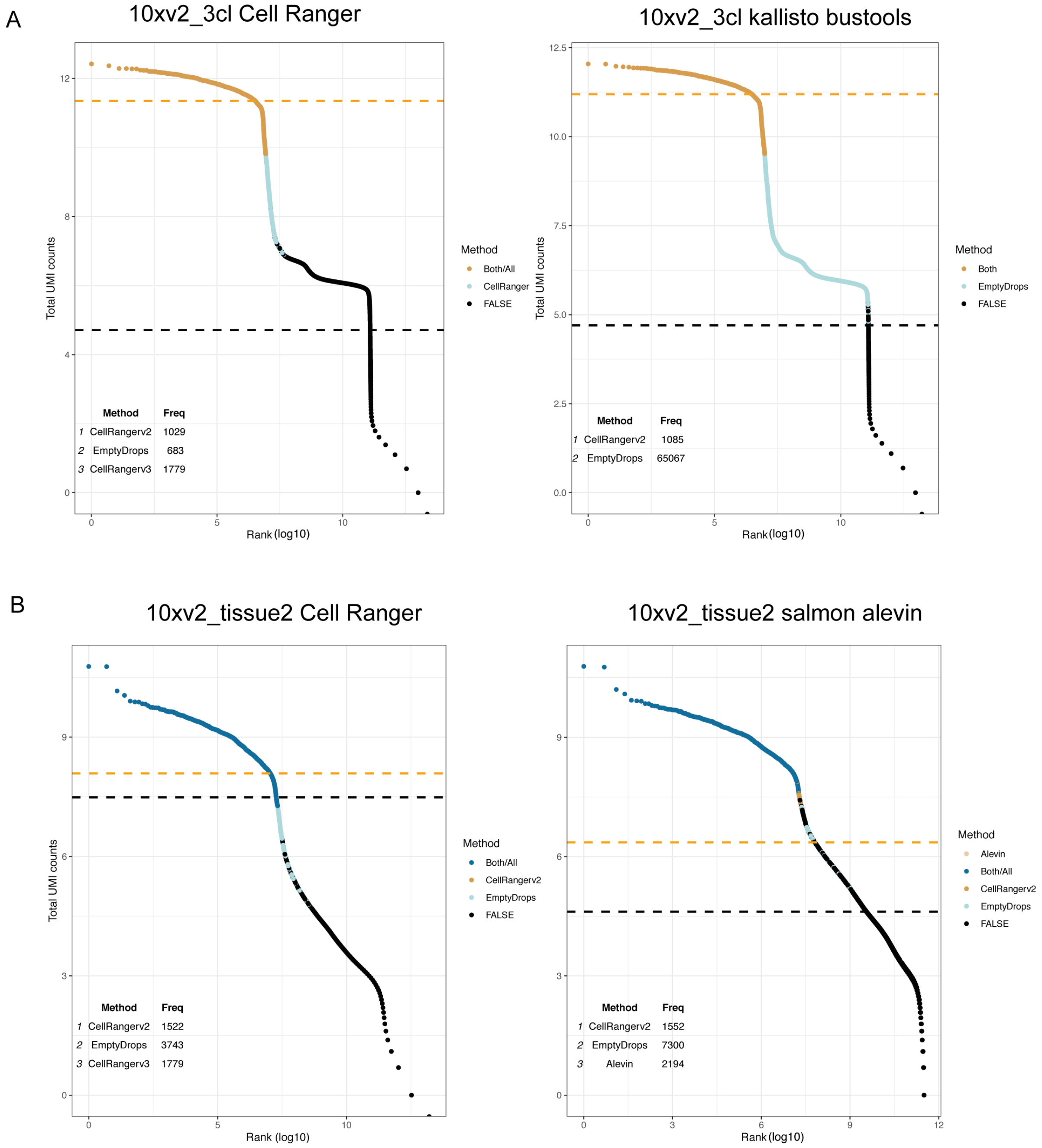


Figure S2, Barcode rank plots generated with *EmptyDrops* and other cell detection methods. A) Results on *Cell Ranger* and *kallisto bustools* generated on 10xv2_3cl datasets. B) Results on *Cell Ranger* and *salmon alevin* generated on 10xv2_tissue2 datasets. The detected knee and inflection points are shown in orange and black accordingly.

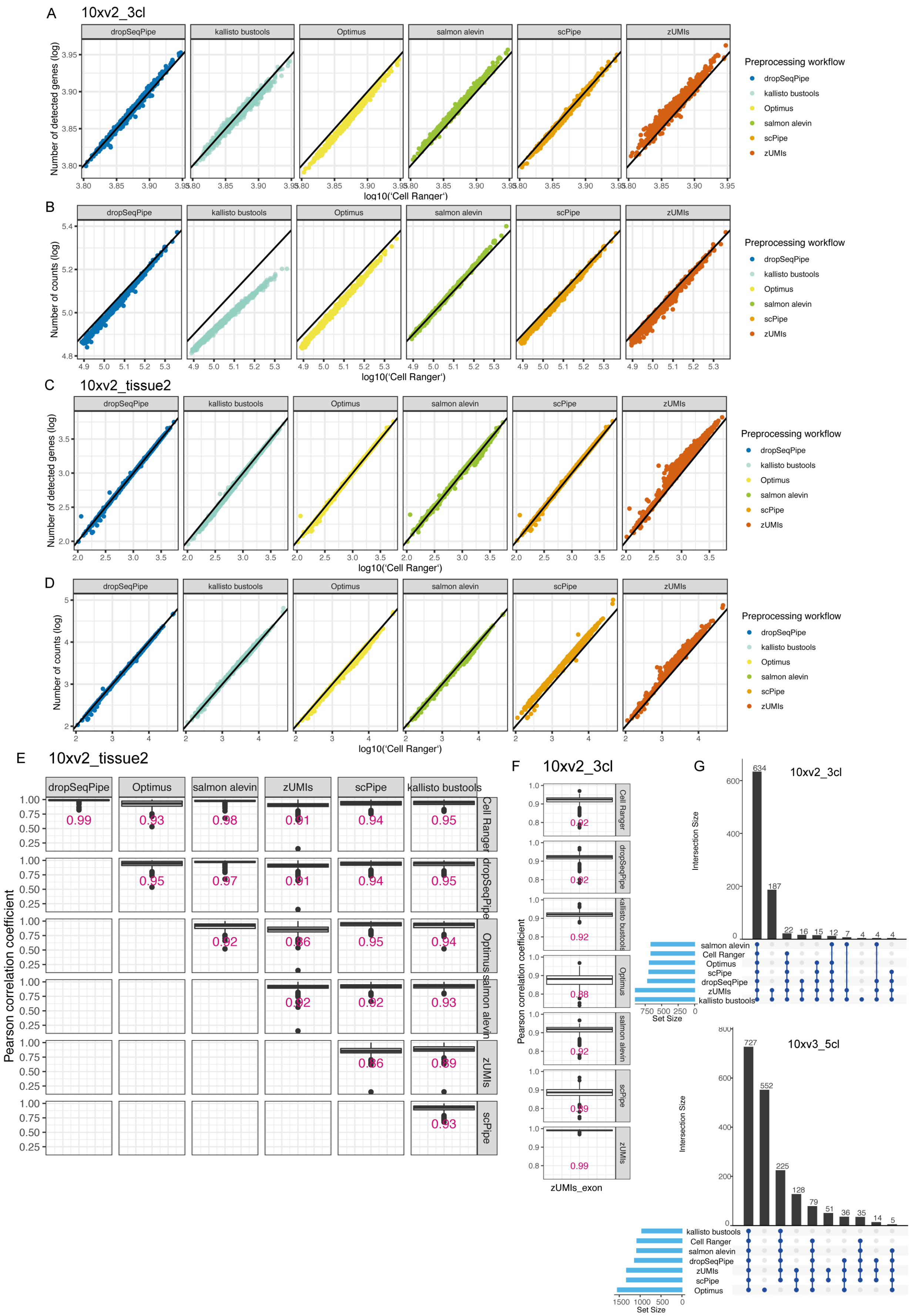


Figure S3, Comparing gene expression quantification across preprocessing workflows on droplet-based datasets. Number of detected genes per cell and total counts per cell of common cells are plotted of listed

preprocessing workflows against *Cell Ranger* accordingly on 10xv2_3cl in A) C) and 10xv2_tissue2 datasets in B) D) (all in log10-scale).
Pearson correlation coefficients are calculated using common genes in individual cells across selected preprocessing workflows on 10xv2_tissue2 datasets, and then plotted in E). The median value of the Pearson correlation coefficients is labelled in the middle of each boxplot. Additionally, Pearson correlation coefficients of *zUMIs*-exon mode with other workflows are shown in F).
The UpSet plots in G) are used to display intersections of retains cells across workflows on 10xv2_3cl, and 10xv3_5cl datasets.

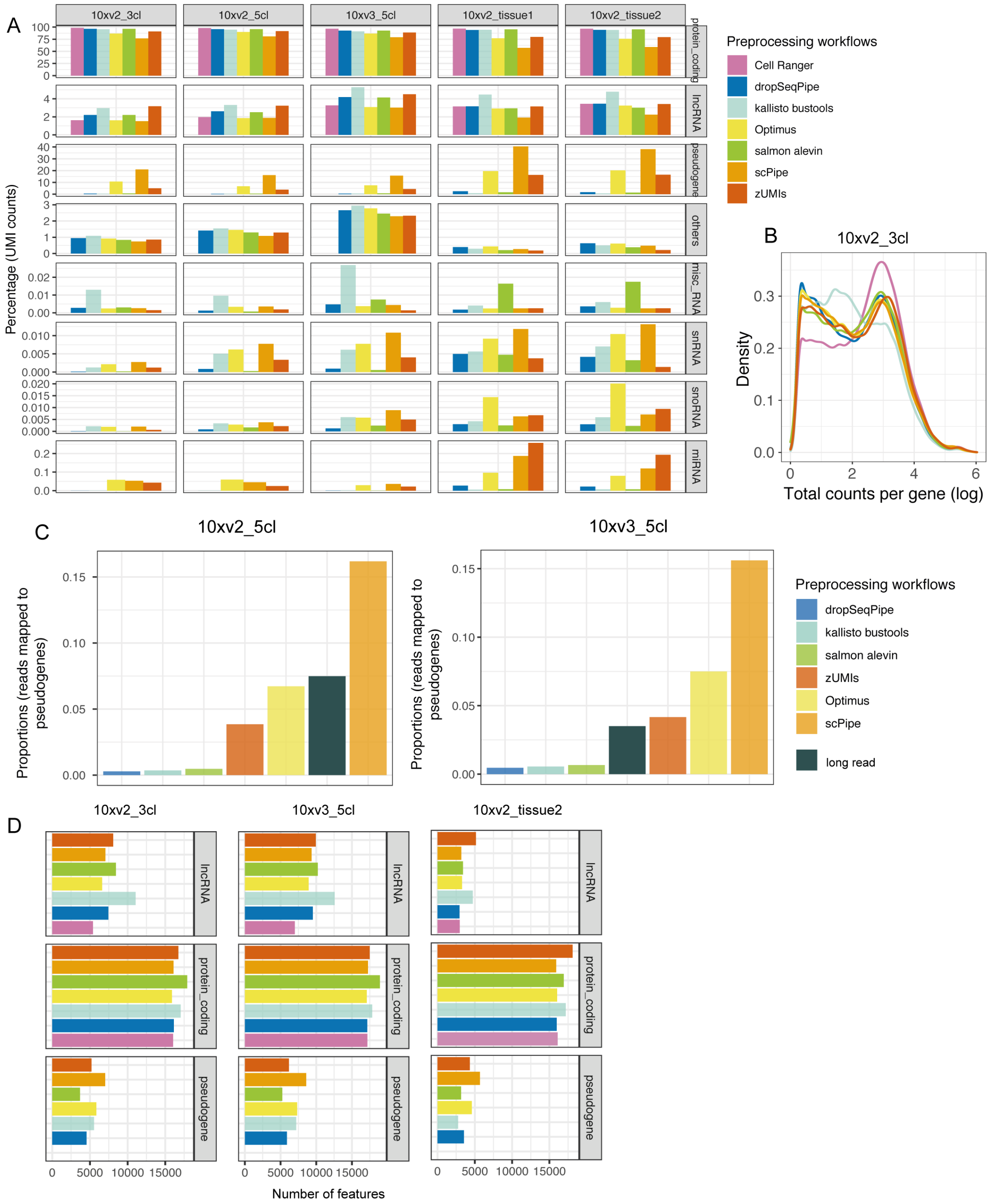


Figure S4, A) Bar plots are used to show proportions of genes of listed gene biotypes delivered across different preprocessing workflows on droplet-based datasets. Gene biotypes of Long non-coding RNAs (lncRNAs), microRNAs (miRNAs), Miscellaneous RNAs (misc-RNAs), protein coding genes, pseudogenes, small nuclear RNAs (snoRNAs), and small nucleolar RNAs (snRNAs) are shown here. Colors represent preprocessing workflows.
The density of total counts per gene on 10xv2_3cl dataset is shown in B).

C), Comparisons of proportions of counts mapped to pseudogenes across selected workflows on short-read sequencing and that obtained from single cell long-read sequencing. Color denotes preprocessing workflows and results from long read data is colored by dark grey.
D), Number of genes of listed gene biotypes delivered by different preprocessing workflows on 10xv2_3cl, 10xv3_5cl, and 10xv2_tissue2 datasets. Only genes of lncRNA, protein coding genes, and pseudogenes were extracted from the raw count matrices and then used in the following evaluation.

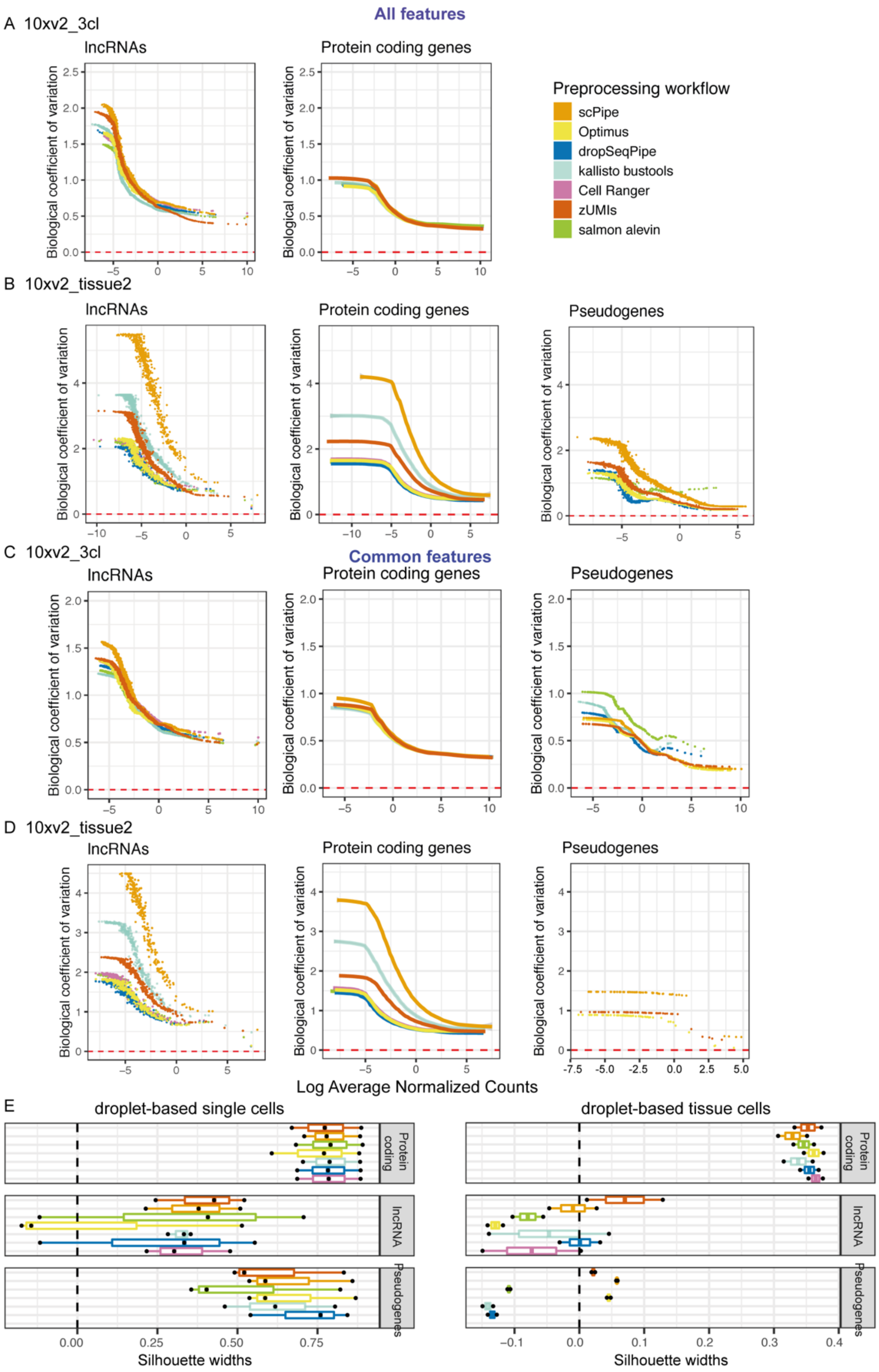


Figure S5, BCV plots of genes with biotypes of lncRNA, protein coding genes and pseudogenes delivered by different preprocessing workflows on A) 10xv2_3cl, and B) 10xv2_tissue2 datasets using common cells. Raw gene counts break down by biotypes are used to calculate biological coefficient of variation (BCV). And then BCVs are plotted against *scran* normalized counts. Colors represent preprocessing workflows. Additionally, BCV are also calculated using common cells and common features and plotted on C) 10xv2_3cl and D) 10xv2_tissue2 datasets. Boxplots of silhouette widths calculated with GLMPCs obtained from different workflows based on known cell types are shown in E) for single cell datasets (10xv2_3cl, 10xv2_5cl, 10xv3_5cl) in the left-hand panel and tissue cell datasets (10xv2_tissue1, 10xv2_tissue2) in the right-hand panel. Silhouette width=0 is shown with a black dashed line.

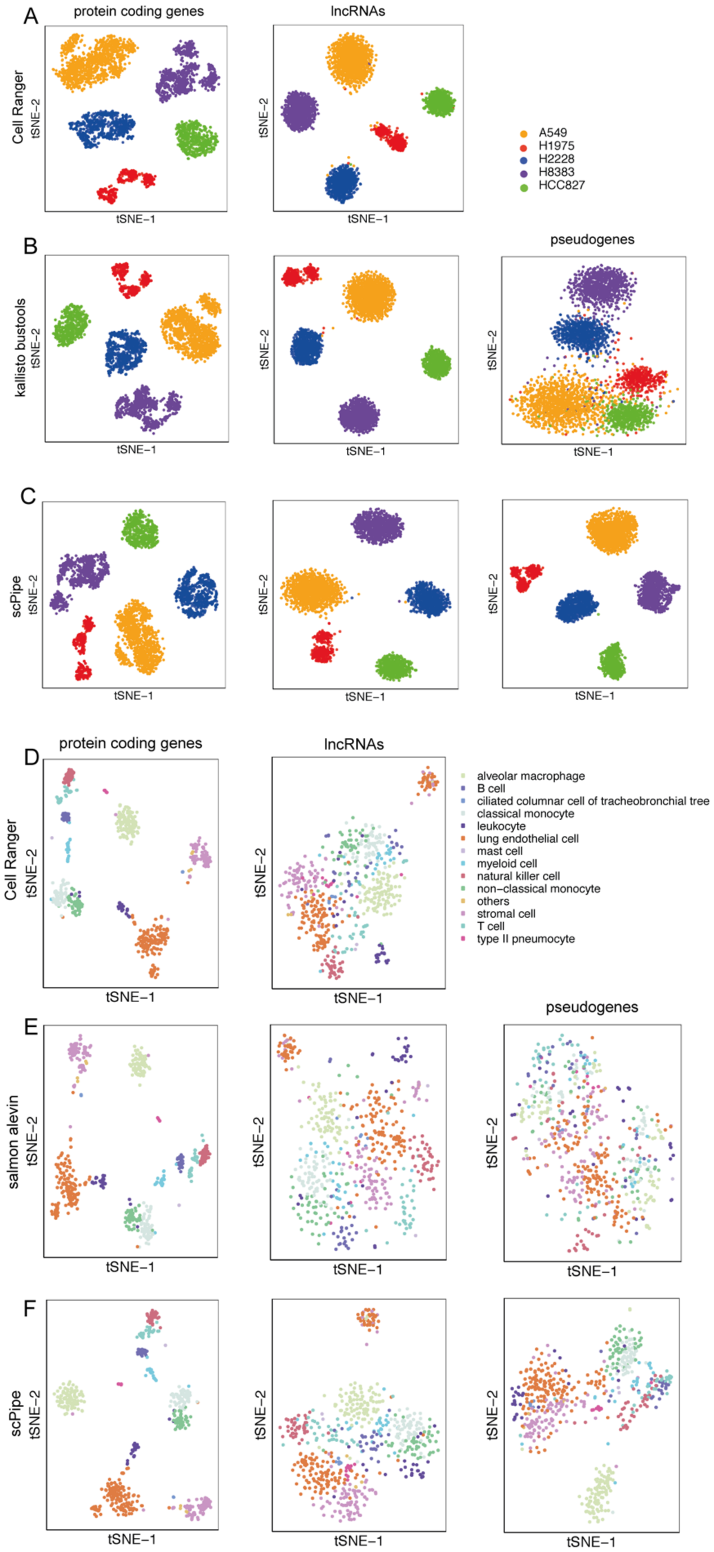


Figure S6, tSNE plots generated with genes of specific gene biotypes on 10xv2_5cl dataset. Protein coding genes, lncRNAs, and pseudogenes are extracted from *scran* normalized counts delivered by A) *Cell Ranger*, B)

*kallisto bustools* and C) *scPipe*, and then visualized with tSNE plots in left, middle and right panel accordingly. Color denotes different cell line cells.
Similarly, on 10xv2_tissue1 dataset, protein coding genes, lncRNAs, and pseudogenes were extracted from *scran* normalized counts delivered by D) *Cell Ranger*, E) *salmon alevin* and F) *scPipe*, and then visualized with tSNE plots in left, middle and right panel accordingly. Color denotes different cell types.

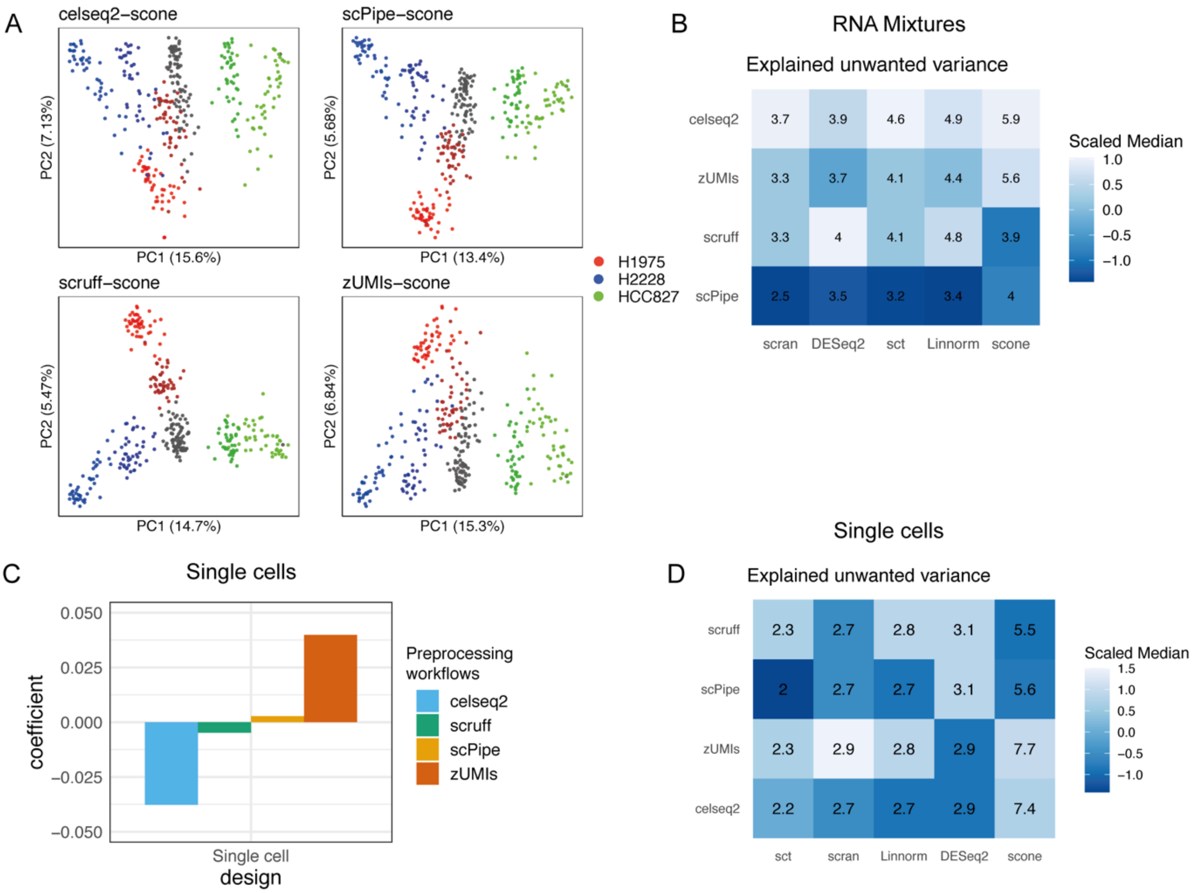


Figure S7, After normalization, on plate-based mixture datasets, PCA plots delivered by combinations of preprocessing workflows and normalization methods are displayed in A). Heatmap of median values of unwanted variance calculated across different preprocessing workflows is shown in B). sct represents *sctransform*.
On plate-based single cells datasets, a linear model is fitted using silhouette widths as dependent variables, with preprocessing workflows as covariates. Coefficients are plotted in C).
Heatmap of median values of unwanted variance calculated across different preprocessing workflows is shown in D).

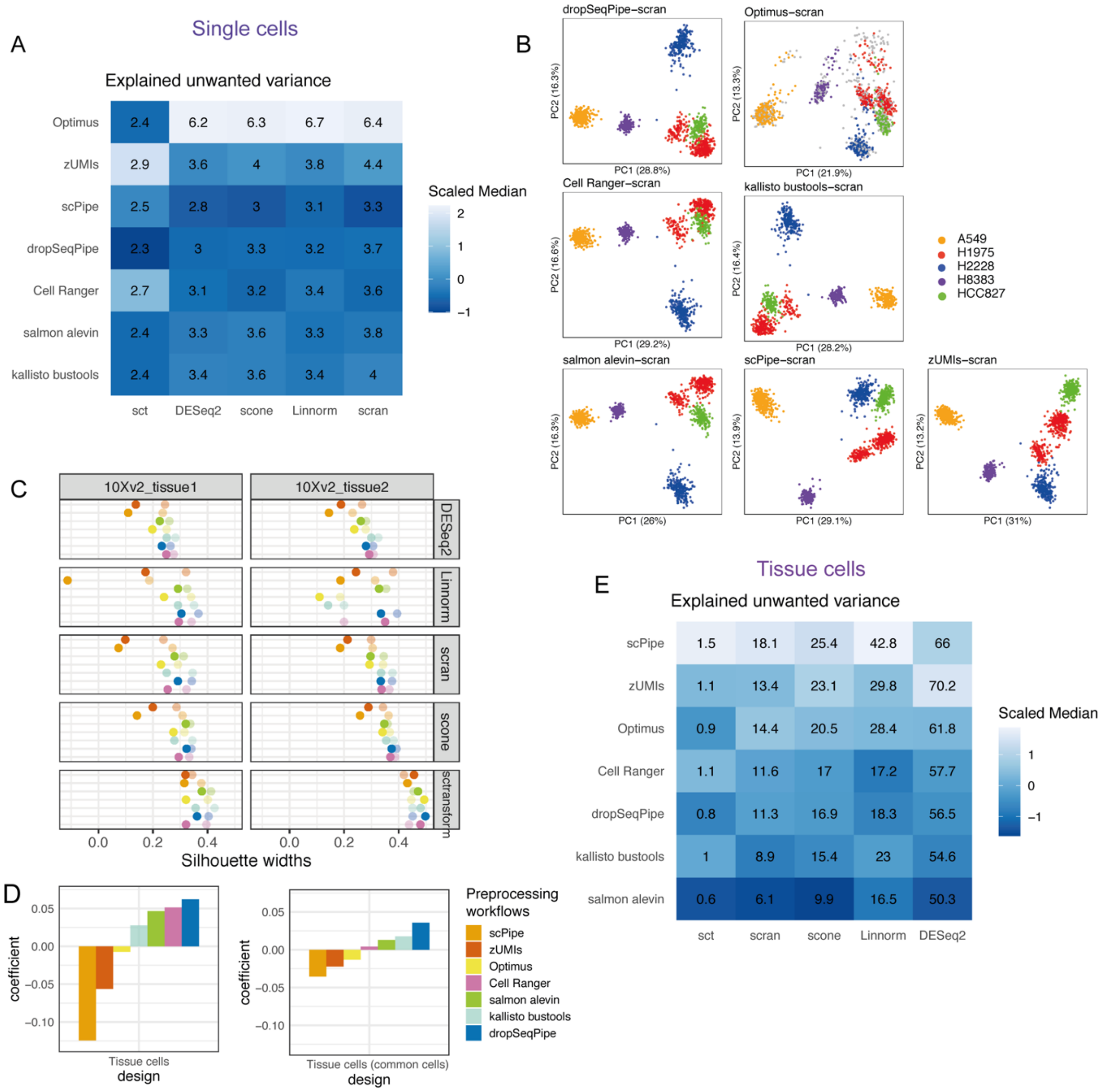


Figure S8, After normalization, on droplet-based single cell datasets, heatmap of median values of explained wanted variance and unwanted variance calculated with different preprocessing workflows is shown in A). PCA plots of scran normalized counts delivered by listed preprocessing workflows on 10xv3_5cl dataset are displayed in B). sct represents *sctransform*.

On droplet-based tissue datasets, the performance of combinations of preprocessing workflows and normalization methods using all cells versus common cells is compared. Silhouette widths calculated using the top 20 PCs based on known cell types are used for comparisons and plotted in C). Here, dots of solid color represent results generated with unfiltered (all) cells, and dots of transparent color represent results generated with common cells across preprocessing workflows.

A linear model is fitted using silhouette widths as dependent variables, with preprocessing workflows as covariates using all cells and common cells separately. Coefficients are plotted in D).

Heatmap of median values of explained unwanted variance calculated with different preprocessing workflows is displayed in E).

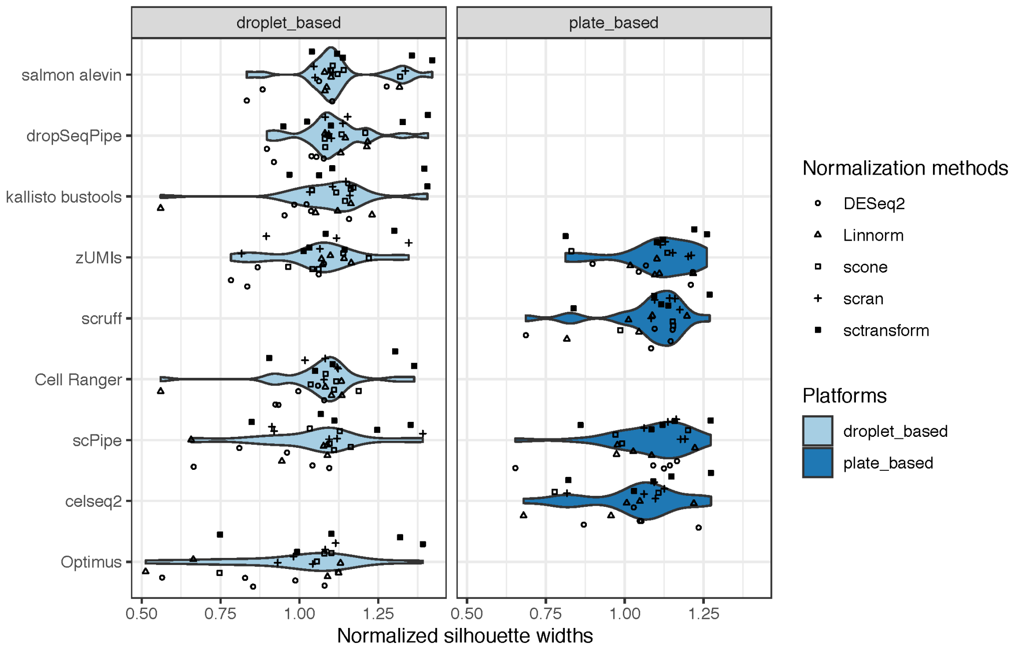


Figure S9, Summary of performance of combinations of preprocessing workflows and normalization methods. From top to bottom, the performance is ordered from the best to the worst according to median normalized silhouette widths. Silhouette widths are calculated based on known cell labels after applying different normalization methods and normalized against the silhouette widths obtained without any normalization. Here, each dot represents a combination. Colors denote different single cell platforms and shapes denote different normalization methods.

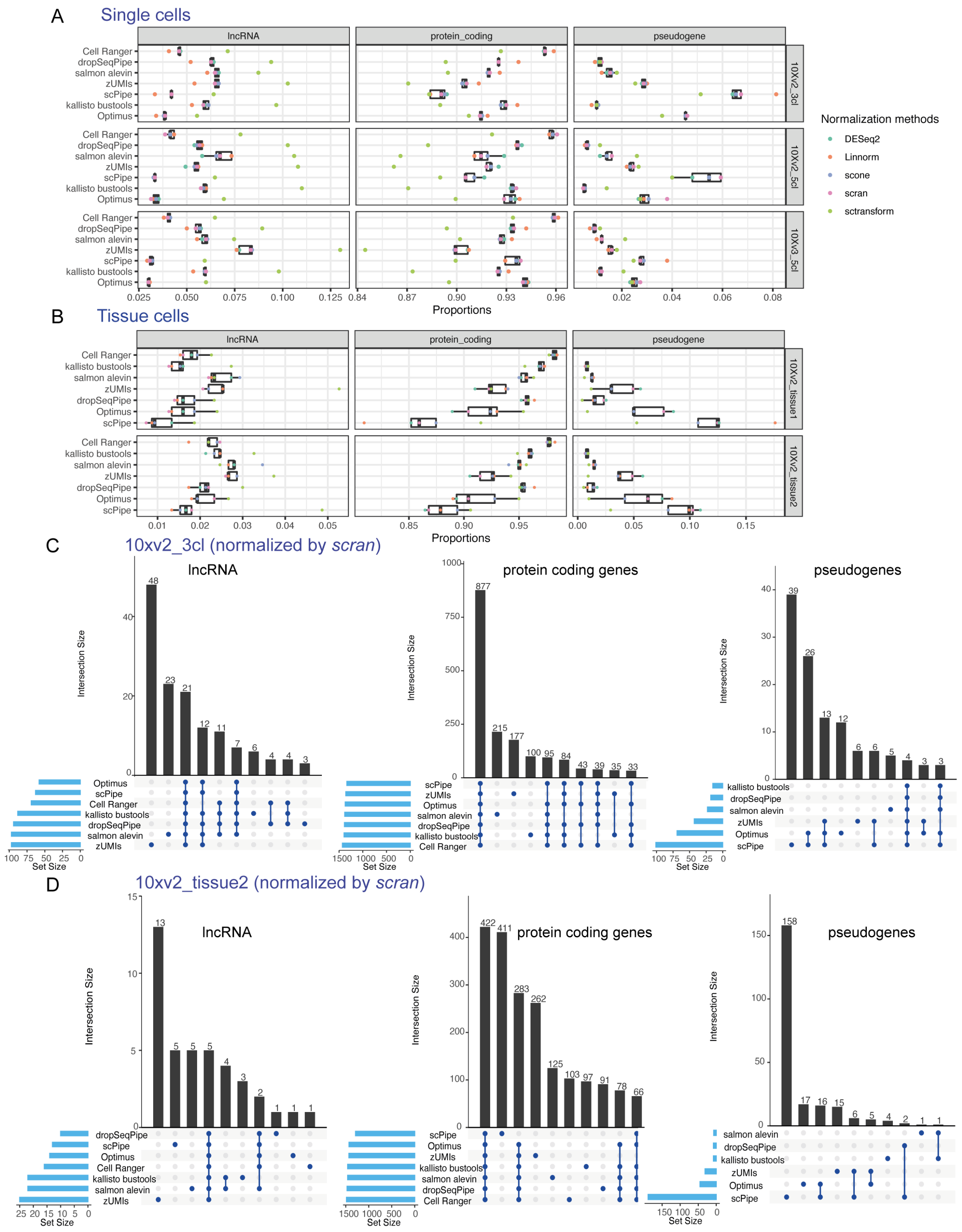


Figure S10, After highly variable gene selection, the percentage of genes of biotypes of lncRNAs, protein coding genes, and pseudogenes in HVGs on droplet-based single cell datasets, including 10xv2_3cl, 10xv2_5cl, and 10xv3_5cl datasets are plotted in A) and on droplet-based tissue datasets, including 10xv2_tissue1 and 10xv2_tissue2 are plotted in B). Color denotes different normalization methods.
Take results normalized by *scran* as examples, intersections across preprocessing workflows of top 1.5k HVGs split by listed gene biotypes on 10xv2_3cl, and 10xv2_tissue1 are shown in C) and D) accordingly using UpSet plots.

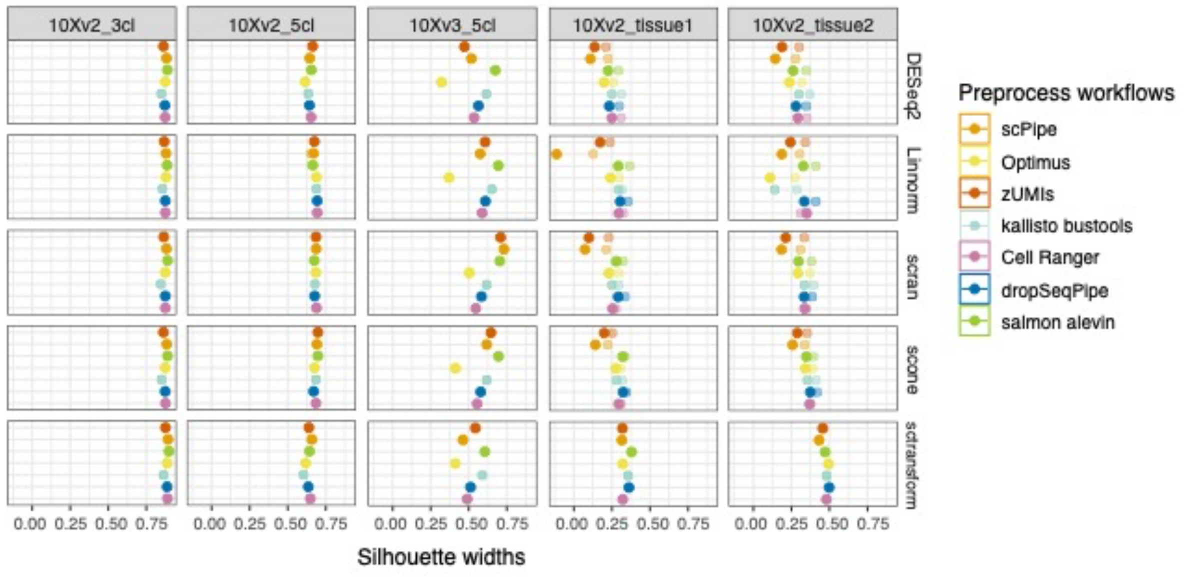


Figure S11, Comparing performance of combinations of preprocessing workflows and normalization methods before and after HVG selection on droplet-based datasets using all cells. Silhouette widths were calculated using PCs by know cell types. Here, dots of solid color represent results before HVG selection and dots of transparent color represent results after HVG selection.

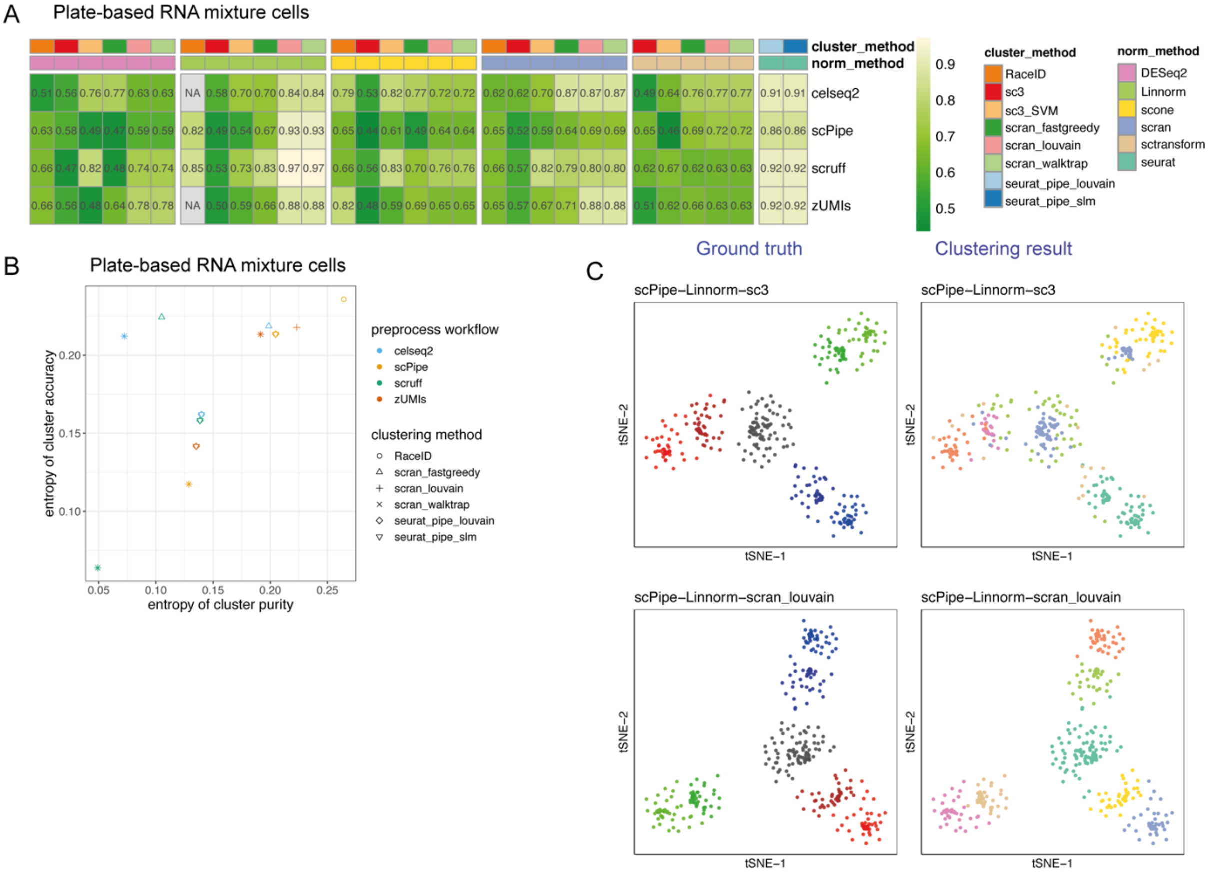


Figure S12, Comparing the impact preprocessing workflows have on clustering results on plate-based RNA mixture datasets. Heatmap of the median value of ARI is shown in A).
ECA versus ECP plot for top 5 combinations delivered by different preprocessing workflows is displayed in B).
Example tSNE plots generated using normalized HVGs delivered by different combinations of preprocessing, normalization, clustering methods are in C). Here, tSNE plots of two combinations are shown. For each

combination, the left tSNE plot is colored by ground truth (known cell labels), and the right tSNE plot is colored by identified clusters.

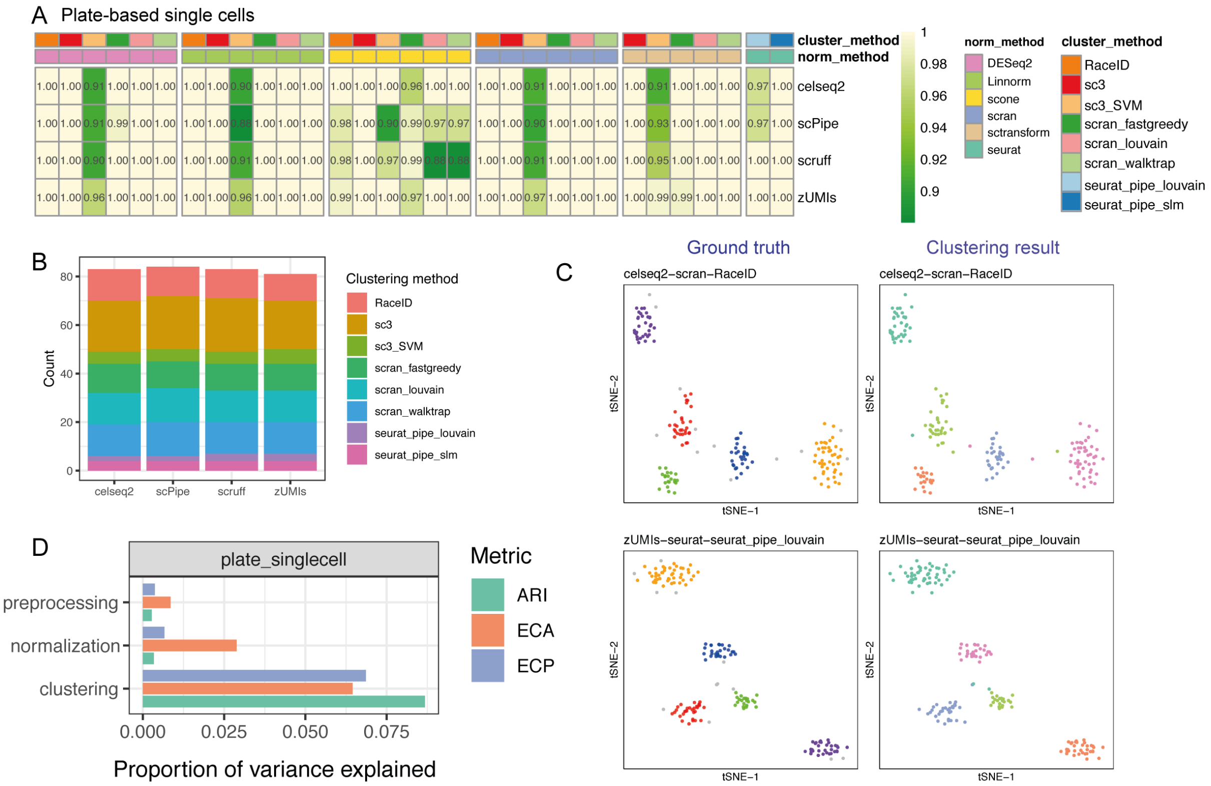


Figure S13, Comparing the impact preprocessing workflows have on clustering results on plate-based single cell datasets. Heatmap of the median value of ARI is shown in A).
B) Number of combinations reached both ECA=0 and ECP=0 across preprocessing workflows on plate-based single cell datasets.
Example tSNE plots generated using normalized HVGs delivered by different combinations of preprocessing, normalization, clustering methods are in C). Here, tSNE plots of two combinations are shown. For each combination, the left tSNE plot is colored by ground truth (known cell labels), and the right tSNE plot is colored by identified clusters.
ANOVA model was used to calculate proportions of variance main analysis steps explained based on clustering evaluation metrics (ARI, ECA and ECP). Results are plotted in D).

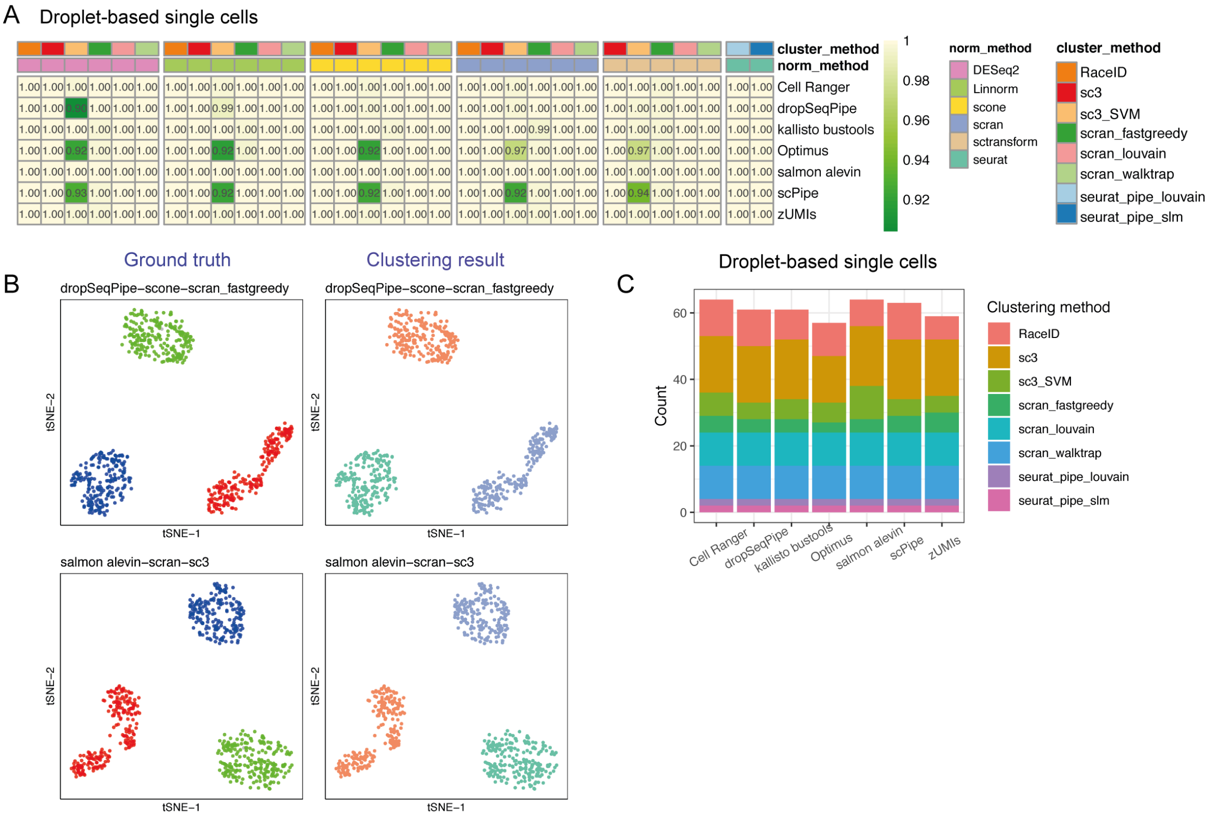


Figure S14, Comparing the impact preprocessing workflows have on clustering results on droplet-based single cell datasets. Heatmap of the median value of ARI is shown in A).

Example tSNE plots generated using normalized HVGs delivered by different combinations of preprocessing, normalization, clustering methods are in B). Here, tSNE plots of two combinations are shown. For each combination, the left tSNE plot is colored by ground truth (known cell labels), and the right tSNE plot is colored by identified clusters.

C) Number of combinations reached both ECA=0 and ECP=0 across preprocessing workflows on droplet-based single cell datasets.

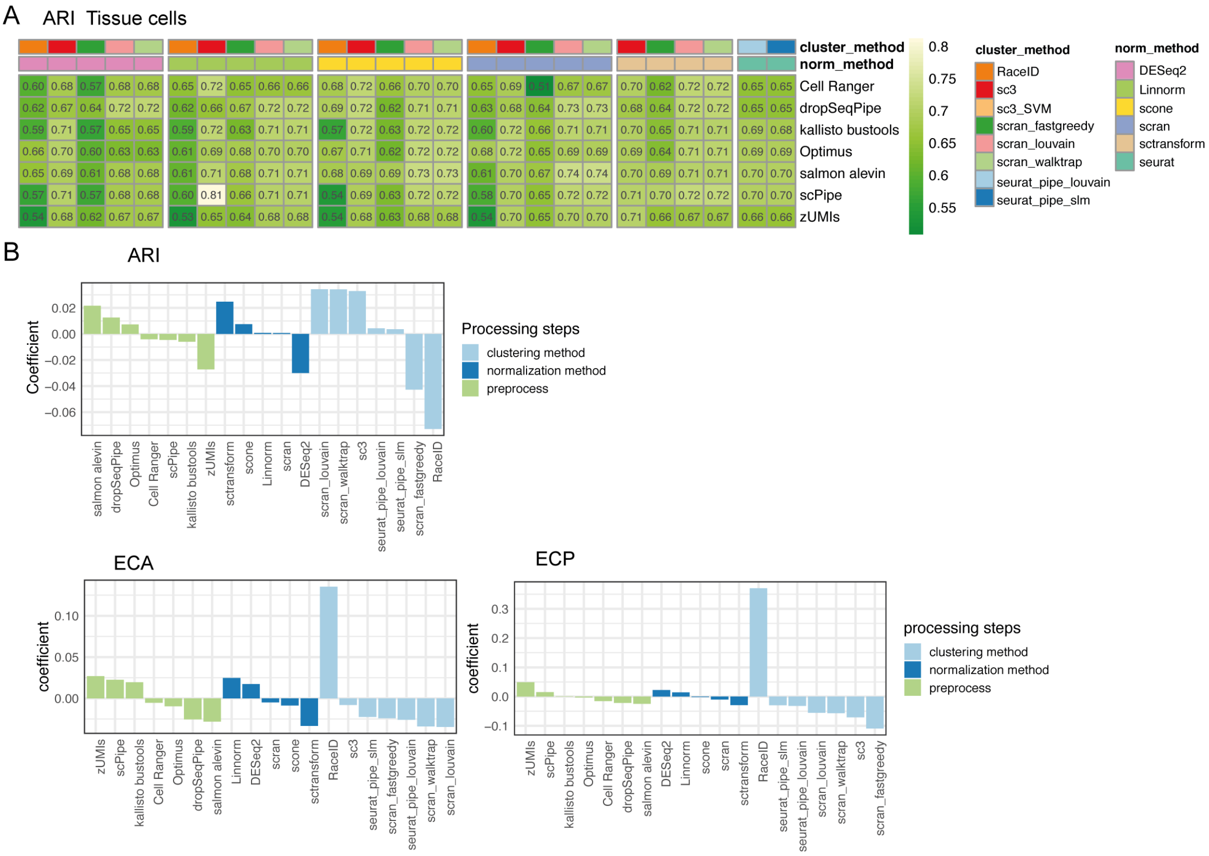


Figure S15, Comparing the impact preprocessing workflows have on clustering results on droplet-based tissue cells datasets.
Heatmap of the median value of ARI using common tissue cells across workflows is shown in A).
Linear models are fitted using evaluation metrics generated with common cells as dependent variables, with experimental design, preprocessing workflows, normalization methods, and clustering methods as covariates. Coefficients calculated with ARI, ECA and ECP are shown in B).
